# mTORC1 activation drives astrocyte reactivity in cortical tubers and brain organoid models of TSC

**DOI:** 10.1101/2025.02.28.640914

**Authors:** Thomas L. Li, John D. Blair, Taesun Yoo, Gerald A. Grant, Dirk Hockemeyer, Brenda E. Porter, Helen S. Bateup

**Author notes:** Equal contribution (Thomas L. Li, John D. Blair).

## Abstract

Tuberous Sclerosis Complex (TSC) is a genetic neurodevelopmental disorder associated with early onset epilepsy, intellectual disability and neuropsychiatric disorders. A hallmark of the disorder is cortical tubers, which are focal malformations of brain development containing dysplastic cells with hyperactive mTORC1 signaling. One barrier to developing therapeutic approaches and understanding the origins of tuber cells is the lack of a model system that recapitulates this pathology. To address this, we established a genetically mosaic cortical organoid system that models a somatic “second-hit” mutation, which is thought to drive the formation of tubers in TSC. With this model, we find that loss of *TSC2* cell-autonomously promotes the differentiation of astrocytes, which exhibit features of a disease-associated reactive state. *TSC2^-/-^* astrocytes have pronounced changes in morphology and upregulation of proteins that are risk factors for neurodegenerative diseases, such as clusterin and APOE. Using multiplexed immunofluorescence in primary tubers from TSC patients, we show that tuber cells with hyperactive mTORC1 activity also express reactive astrocyte proteins, and we identify a unique population of cells with expression profiles that match those observed in organoids. Together, this work reveals that reactive astrogliosis is a primary feature of TSC that arises early in cortical development. Dysfunctional glia are therefore poised to be drivers of pathophysiology, nominating a potential therapeutic target for treating TSC and related mTORopathies.

## Introduction

Tuberous Sclerosis Complex (TSC) is a multisystem developmental disorder caused by mutations in *TSC1* or *TSC2^1^*. TSC affects approximately 1:6000 live births^2^, causing hamartomas in multiple organs and a constellation of neurological and psychiatric conditions, including early-onset epilepsy^3^. Focal brain malformations, known as cortical tubers, are a hallmark of TSC. Tubers are observable by MRI and are composed of dysplastic neurons and glia^4^. A subset of tubers and surrounding regions can become seizure foci, and surgical removal can lead to seizure freedom in individuals with intractable epilepsy^5^. Micro-tubers, regions containing clusters of dysplastic cells, have also been observed throughout the brain in post-mortem samples^6^.

The developmental origin of tuber cells and the molecular changes that lead to their abnormal development are open questions. On a cellular level, tubers contain dysmorphic neurons, dysplastic and gliotic astrocytes, and giant or balloon cells within a background of normal-appearing cells^4^. Gene expression analysis of tuber tissue has revealed an increase in immune response genes as well as astrocyte-enriched genes^7,8^, but these analyses are confounded by the heterogeneity of cell types within tubers^4,9,10^. While much research has focused on neuronal mechanisms in TSC, some studies have suggested that the primary pathological cells within tubers may be astrocytes^11^. However, since tuber resections are performed in individuals with intractable epilepsy, it has been unclear whether glial changes are a cause or consequence of ongoing seizures. An additional challenge in understanding the pathophysiological mechanisms of tubers is lack of a clear cell type signature, as dysplastic cells can express progenitor cell markers as well as neuronal and glial proteins^12-15^.

A prevailing hypothesis for how *TSC1/*2 mutations lead to tubers is through a “second-hit” mutation. In this model, patients have a germline heterozygous mutation and acquire a somatic mutation in the functional allele in a subset of progenitor cells causing loss of heterozygosity (LOH)^16^. *TSC1* and *TSC2* encode proteins forming a multimeric complex that negatively regulates mTOR complex 1 (mTORC1)^17^, a key signaling node that balances cellular anabolic and catabolic processes^18^. Loss of function of either TSC1 or TSC2 results in mTORC1 hyperactivity^19^. Therefore, while second-hit progenitor cells survive and proliferate, they are impaired in their differentiation and development, giving rise to focal regions of abnormal cells. The second-hit model explains the localized, stochastic, and variable nature of tubers, as well as the observation that only a subset of cells within tubers show high mTORC1 activity^4,6^. Second-hit mutations are frequently found in TSC-related hamartomas and have been identified in some resected patient tubers^7,20-22^. Independent of TSC, the majority of focal cortical dysplasia type II cases, which have overlapping histological and clinical features with TSC, are explained by somatic mutations in the mTOR pathway, including second-hit mutations^23,24^. However, not all tubers have an identified second-hit mutation, and it has been suggested that tuber-like cells may arise from heterozygous cells in some contexts^25^.

Here, we use a human brain organoid model that recapitulates the second-hit mechanism to uncover the developmental and molecular alterations in *TSC2^-/-^* cells. Our previous work showed that loss of *TSC2* in organoid models leads to mTORC1 hyperactivity and the generation of hypertrophic glial-lineage cells^26^. In this study, we use single cell-transcriptomics, cell-type isolation and high-throughput image analysis across multiple stem cell lines to demonstrate that these glial-lineage cells are reactive astrocytes. We show that this reactive state emerges cell-autonomously, in contrast to other situations in which astrocytes are activated in response to signals from other brain cells^27^. We identify novel, differentially expressed genes in tuber-like cells in organoids, including several that are known risk factors for neurodegenerative diseases including *APOE^28^* and *CLU^29^*. Lastly, using a cyclic immunostaining approach, we identify a unique population of cells in patient tubers that have hyperactive mTORC1 signaling and express the key astrocyte genes identified in the organoid model.

This work shows that a second-hit mutation in *TSC2* can generate the dysmorphic cells seen in tubers. The fact that the glial changes observed in patient tissue are recapitulated in developing cortical organoids provides evidence that glial dysfunction is an early driver of pathophysiology in TSC, rather than just a response to ongoing seizures. Together our work establishes reactive astrocytes as relevant therapeutic targets for TSC and related mTORopathies.

## Results

### Two-hit *TSC2^-/-^* cells generate transcriptionally activated astrocytes in forebrain organoids

To understand the developmental processes leading to tuber cell formation, we performed single-cell RNA sequencing (scRNAseq) of a genetically mosaic human brain organoid model that recapitulates a second-hit mutation^26^. These cells have a loss-of-function mutation in one allele of *TSC2* (exon 5 deletion), and a conditional allele that can be rendered non-functional by Cre recombinase (Extended Data Fig. 1a-d). Second-hit cells are identified by red fluorescence via a Cre-dependent tdTomato knock-in reporter (Fig. 1a, Extended Data Fig. 1c). We differentiated *TSC2^c/-;LSL-TdTom^* human embryonic stem cells (hESCs) into forebrain organoids using a protocol^30,31^ that recapitulates developmental transitions including the proliferation of neural progenitor cells, neurogenesis, and subsequently gliogenesis and cellular maturation. Cre recombinase was delivered via lentiviral transduction at day 8 to delete *TSC2* in a subset of cells in the organoid (Fig. 1b). We performed fluorescence-activated cell sorting (FACS) and scRNA-seq of *TSC2^c/-;LSL-TdTom^* organoids at day 50 (neurogenesis), day 120 (early gliogenesis), and day 220 (cell maturation) (Fig. 1b). Organoids from three independent differentiations were collected at each time point, with each organoid contributing both heterozygous (*TSC2^c/-^*; tdTomato-negative) and homozygous (*TSC2^-/-^*; tdTomato-positive) cells.

**Figure 1:**
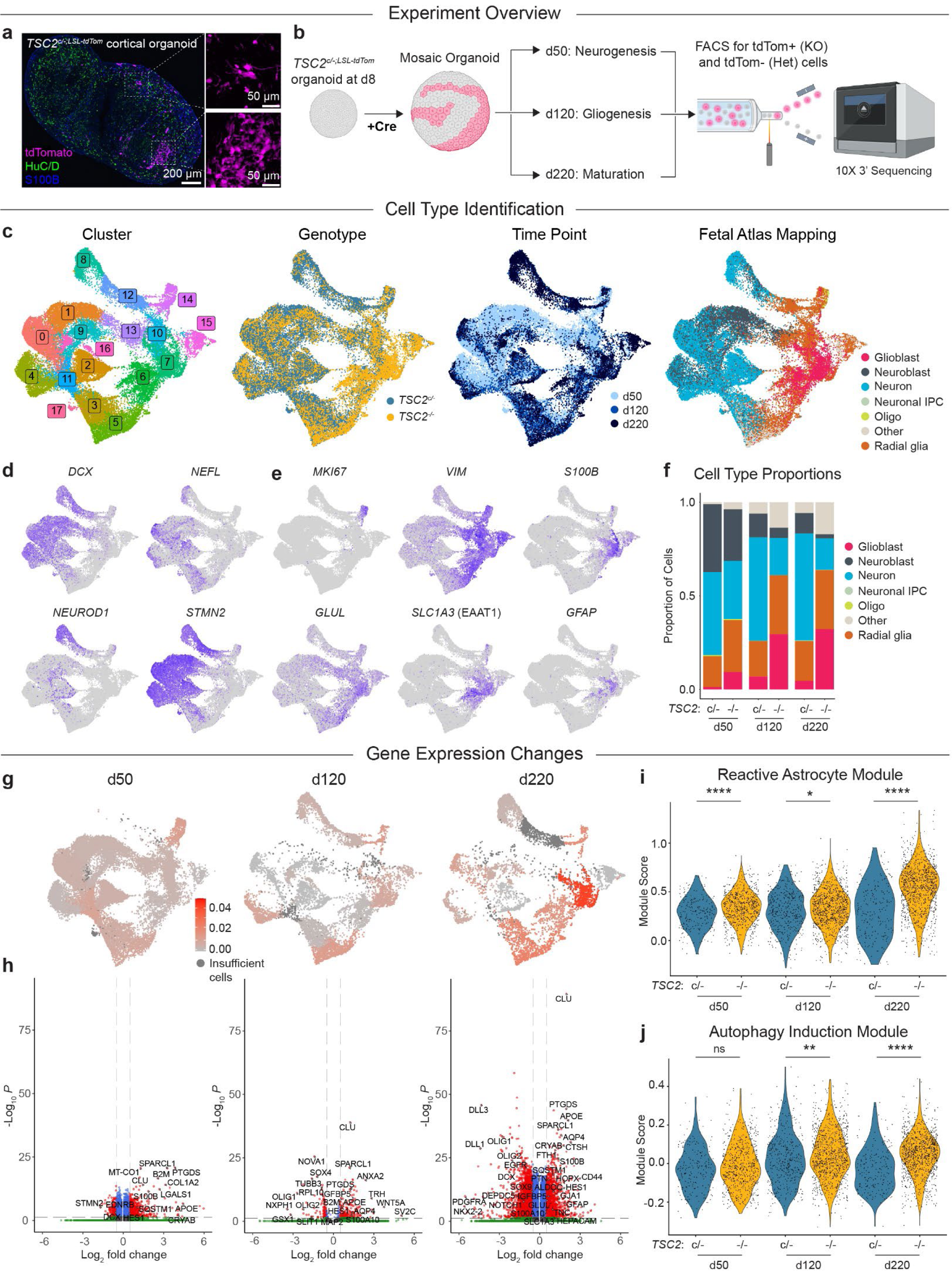
Single-cell RNA sequencing of WIBR3 *TSC2^c/-;LSL-TdTom^* brain organoids. **a,** Example image of an immunostained WIBR3 *TSC2^c/-;LSL-TdTom^* organoid showing tdTomato-positive *TSC2^-/-^* cells. HuC/D labels neurons in green and S100β labels glial cells in blue. **b,** Schematic of the experimental design. Brain organoids were generated from WIBR3 *TSC2^c/-;LSL-TdTom^* hESCs and exposed to Cre lentivirus at day 8. Organoids were cultured until day 50, 120, or 220, at which time they were dissociated, separated by FACS, and processed for 10x scRNA-seq. **c,** UMAP plots of scRNA-seq results, grouping cells by unbiased clustering, genotype, time point, and mapping to a fetal brain atlas. **d,** Feature plots of selected neuronal genes. **e,** Feature plots of selected astrocyte and neural progenitor genes. **f**, Cell type proportions, as assayed by atlas mapping, divided by genotype and time point. **g**, Cluster-based normalized expression distances between *TSC2^-/-^* and *TSC2^c/-^* cells. Expression distances were calculated between the two genotypes within each cluster. **h**, Volcano plots of differential gene expression between *TSC2^-/-^* and *TSC2^c/-^* cells in cluster 7, an astrocyte cluster, at day 50, 120, and 220. Genes more highly expressed in *TSC2^-/-^* cells have a positive Log_2_ fold change, and genes more highly expressed in *TSC2^c/-^* cells have a negative Log_2_ fold change. **i,** Violin plots by genotype and time point displaying scores for a reactive astrocyte gene module within glioblast-mapped cells. Violins show overall distributions and dots represent values for individual cells. **j,** Module scores per cell for an autophagy induction gene module within glioblast-mapped cells. See Supplemental Table 10 for statistics and sample sizes.

After processing, 39,539 cells were mapped to a fetal brain tissue atlas^32^ (Fig. 1c). Cell type mapping was confirmed with feature plots of key neuronal genes including *DCX, NEFL, NEUROD1,* and *STMN2* (Fig. 1d), glial genes such as *VIM, S100B, GLUL, SLC1A3,* and *GFAP,* and the proliferation marker *MKI67* (Fig. 1e). Analysis of cell type proportions revealed that while *TSC2^-/-^* cells could be mapped onto known cell types, they were heavily biased toward radial glial and glioblast fates at all timepoints (Fig. 1f). Specifically, *TSC2^-/-^* cells generated more glioblasts, which includes astrocyte progenitor cells and astrocytes, compared to *TSC2^c/-^* cells in the same organoid as early as day 50, with this bias growing more prominent over time (Fig. 1f). While previous work relied on selected marker genes^26^, this transcriptomic mapping shows that *TSC2^-/-^* cells have the gene expression profile of astrocytes.

*TSC2^-/-^* progenitors not only generated a greater proportion of glioblasts, but these cells had the greatest transcriptional changes as a result of TSC2 loss. We performed expression distance calculations^33^ between genotypes in each cluster at each time point (Fig. 1g) and found the greatest transcriptional changes in cluster 7, an astrocyte cluster defined by expression of *S100B, GLAST* and *GLUL*. Compared to *TSC2^c/-^* cells, *TSC2^-/-^* cells had reduced expression of neuronal genes (*STMN2* and *DCX*) and oligodendrocyte genes (*PDGFRA, OLIG1* and *OLIG2*), with striking upregulation of astrocyte genes including *CLU, S100B, PTGDS, APOE, CRYAB, AQP4,* and *GFAP* (Fig. 1h). Both the number of differentially expressed (DE) genes and the magnitude of the differential expression increased at later time points (Fig. 1h, Supplemental Table 1)

We noted that many of the differentially expressed (DE) genes in *TSC2^-/-^* astrocytes were previously shown to be associated with a disease-associated “reactive” state^34^. To investigate this further, we calculated a reactive astrocyte module score for each glioblast-mapped cell, based on the expression of genes that are upregulated when astrocytes enter an activated state^34^ (Supplemental Table 2). Compared to *TSC2^c/-^* astrocytes, *TSC2^-/-^* astrocytes had significantly higher expression of this module at all time points (Fig. 1i). Since this analysis compares *TSC2^c/-^* and *TSC2^-/-^* cells from the same organoids, these differences cannot be explained by variation in culture conditions, environment, or signals from other cells. Therefore, this result indicates that loss of *TSC2* drives cell autonomous astrocyte activation.

In addition to astrocytic genes, autophagy genes were upregulated in *TSC2^-/-^* astrocytes, including *SQSTM1*, which encodes p62. A module score related to the initiation of autophagy^35^ showed that *TSC2^-/-^* cells had increased expression of this module (Fig. 1j, Supplemental Table 2). Since loss of TSC1/2 and activation of mTORC1 is typically associated with decreased functional autophagy^36^, this may reflect a compensatory response. To examine molecular pathway changes in an unbiased manner, we performed Single Cell Pathway Analysis (SCPA)^37^ to assess changes in gene ontology modules between *TSC2^-/-^* and *TSC2^c/-^* cells on a cluster-by-cluster basis. In both neurons and glia, we found that the most highly enriched pathways were related to translation, metabolism, and cellular stress response (Extended Data Fig. 2a,b, Supplemental Table 3).

It has been suggested that tubers might arise from cells with a heterozygous loss of *TSC1* or *TSC2*^25^, however other studies have reported that heterozygous cells do not exhibit the full characteristics of tuber cells^38,39^. To address this, we generated a *TSC2^c/+;LSL-TdTom^* hESC line, which contains a conditional loss-of-function allele and tdTomato reporter but with a functional second allele. In this model, Cre recombinase generates a “single-hit”, resulting in loss of one copy of *TSC2*. TdTomato-positive cells in *TSC2^c/+;LSL-TdTom^* organoids did not have dysmorphic morphology, a glial differentiation bias, or increased phosphorylation of the mTORC1 pathway target S6 (p-S6) (Extended Data Fig. 3a,b). We performed scRNAseq of *TSC2^c/+;LSL-TdTom^* organoids at days 50 and 120 and did not find a substantial cell type bias or major changes in gene expression (Extended Data Fig. 3c-h, Supplemental Tables 4,5). While *TSC2^c/-^* glioblasts did show modest activation of the reactive astrocyte gene module (Extended Data Fig. 3i), this was less pronounced than in *TSC2^-/-^* cells. These data show that biallelic loss of *TSC2* is required for the production of dysmorphic, tuber-like cells in our brain organoid model.

### *TSC2^-/-^* astrocyte phenotypes are robust across different genetic backgrounds

Variations in the genetic background of stem cell lines can affect how disease-associated mutations influence neural differentiation and development^40^. The WIBR3 *TSC2^c/-;LSL-TdTom^* line is a female hESC line^41^. To test the robustness of our findings, we performed similar scRNAseq experiments in a male induced pluripotent cell (hiPSC) line derived from BJ fibroblasts^42^, which we engineered to carry the conditional *TSC2* knockout machinery (Extended Data Fig. 1a-d). These stem cells were differentiated into brain organoids, exposed to Cre at day 8, and scRNAseq on five differentiation batches was performed on day 140 (Fig. 2a-d). This dataset was analyzed separately to preserve the internally-controlled experimental design. We noted that organoids derived from BJ hiPSCs exhibited stronger mapping to the dorsal pallium, with neuronal clusters corresponding to populations of cortical excitatory and inhibitory neurons (Extended Data Fig. 4a-d).

**Figure 2:**
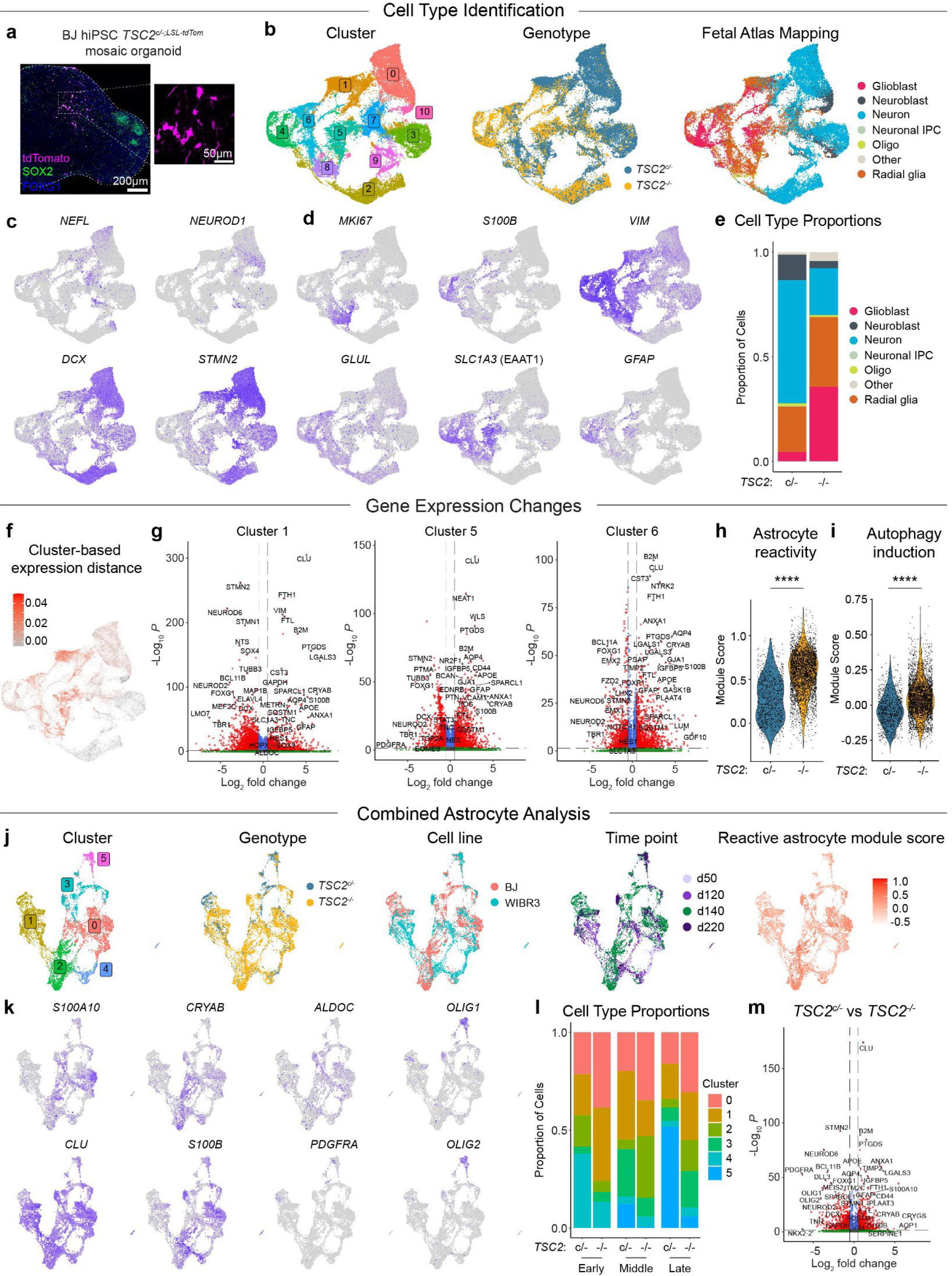
Single-cell RNA sequencing of BJ *TSC2^c/-;LSL-TdTom^* brain organoids and combined astrocyte analysis. **a,** Example image of an immunostained BJ *TSC2^c/-;LSL-TdTom^* organoid showing tdTomato-positive *TSC2^-/-^* cells. SOX2 (green) and FOXG1 (blue) label forebrain progenitor cells. **b,** UMAP plots of scRNA-seq results from BJ *TSC2^c/-;LSL-TdTom^* brain organoids at day 140, grouping cells by unbiased clustering, genotype, and mapping to a fetal brain atlas. **c,** Feature plots of selected neuronal genes. **d,** Feature plots of selected astrocyte and progenitor genes. **e**, Cell type proportions, as assayed by mapping to a fetal brain atlas, divided by genotype. **f**, Cluster-based normalized expression distances between *TSC2^-/-^* and *TSC2^c/-^* cells. Expression distances were calculated between the two genotypes within each cluster. **g,** Volcano plots of differential gene expression between *TSC2^-/-^* and *TSC2^c/-^* cells in astrocyte clusters 1, 5, and 6. Genes more highly expressed in *TSC2^-/-^* cells have a positive Log_2_ fold change, and genes more highly expressed in *TSC2^c/-^* cells have a negative Log_2_ fold change. **h,** Violin plots displaying scores for a reactive astrocyte gene module. Violins show overall distributions and dots represent values for individual cells. **i,** Module scores per cell for an autophagy induction gene module. **j,** UMAP plots of glioblast-mapped cells from the WIBR3 and BJ organoids, divided by unbiased clustering, genotype, cell line, time point, and reactive astrocyte module score. **k**, Feature plots of selected astrocyte and oligodendrocyte precursor cell genes. **l**, Cluster proportions, divided by time point and genotype. The “early” time point consists of cells from day 50 organoids, the “middle” time point consists of cells from day 120 and 140 organoids, and the “late” time point consists of cells from day 220 organoids. **m**, Volcano plot of differential gene expression between *TSC2^-/-^* and *TSC2^c/-^* glioblasts across both hPSC lines at day 120-140. Genes more highly expressed in *TSC2^-/-^* cells have a positive Log_2_ fold change, and genes more highly expressed in *TSC2^c/-^* cells have a negative Log_2_ fold change. See Supplemental Table 10 for statistics and sample sizes.

In BJ *TSC2^c/-;LSL-TdTom^* organoids, we again observed a pronounced astrocyte differentiation bias, the generation of cell-autonomously reactive astrocytes, and activation of the autophagy induction module (Fig. 2e-i, Supplemental Tables 6,7, and Extended Data Fig 2c,d). To test whether *TSC2^-/-^* glioblasts represented a distinct cell type or state, we performed a combined analysis of glioblasts from the WIBR3 and BJ *TSC2^c/-;LSL-TdTom^* lines (Fig. 2j). Reclustering of all glioblasts revealed that *TSC2^-/-^* cells were more highly represented in cluster 0, which was defined by reactivity genes such as *S100A10* and *CRYAB* (Fig. 2k,l). *TSC2^c/-^* cells were more represented in cluster 5, which expressed oligodendrocyte precursor genes such as *PDGFRA*, *OLIG1,* and *OLIG2* (Fig. 2k,l). This shift demonstrates that *TSC2^-/-^* cells preferentially differentiate into astrocytes and not other glial fates such as oligodendrocytes. A combined DE analysis on the day 120-140 cells from both lines showed that the most upregulated genes in *TSC2^-/-^* astrocytes were *CLU, B2M, PTGDS, APOE, S100A10, GFAP, CRYAB,* and *S100B*, with a notable reduction in neuronal and oligodendrocyte-lineage genes (Fig. 2m, Supplemental Table 8).

Previous studies have suggested that tuber cells may be in a progenitor-like or immature state due to their expression of progenitor markers such as nestin and vimentin^16,43^. Organoid-derived astrocytes are known to recapitulate maturity-related changes in astrocyte gene expression through fetal and postnatal development^44,45^. To assess how loss of TSC2 affected cell maturity, we calculated immature and mature astrocyte gene module scores for glioblasts in both stem cell lines (Supplemental Table 2). We found that *TSC2^-/-^* glioblasts had higher maturity scores and lower immaturity scores than corresponding *TSC2^c/-^* glioblasts at each time point (Extended Data Fig. 4e-h). This indicates that at least in organoids, *TSC2^-/-^* astrocytes are not arrested in an immature state.

In summary, our scRNAseq analysis across multiple genetic backgrounds demonstrates that *TSC2^-/-^* cells produce disproportionately more astrocytes than *TSC2^c/-^* cells in the same organoid, and that these astrocytes express higher levels of genes associated with a reactive state. Importantly, these changes are cell-autonomous and triggered by loss of *TSC2* alone, in the absence of inflammatory signals from microglia or other cells.

### *TSC2^-/-^* astrocytes are hypertrophic and have altered protein expression

We hypothesized that the reactive astrocytes generated by *TSC2^-/-^* cells in organoids may represent the dysmorphic glia seen in tubers. To facilitate analysis of protein level changes, we generated isogenic *TSC2^-/-^* and *TSC2^+/+^* hiPSCs, in which all cells have the same genotype, on two different genetic backgrounds (8858 and 8119 hiPSCs^31^) (Fig. 3a). *TSC2^+/+^* hiPSCs went through the gene editing pipeline but remained wild-type at the *TSC2* locus to control for any effects of Cas9 exposure and clonal isolation. *TSC2^-/-^* organoids generated from these stem cells had the expected increases in mTORC1 activity, as well as changes in astrocyte genes at the mRNA and protein levels (Fig. 3b-g, Extended Data Fig. 5). Using an automated analysis to identify cells positive for cell type markers, we confirmed that *TSC2^-/-^* organoids had a significantly higher glia to neuron ratio across both hiPSC lines (Fig. 3h-j).

**Figure 3:**
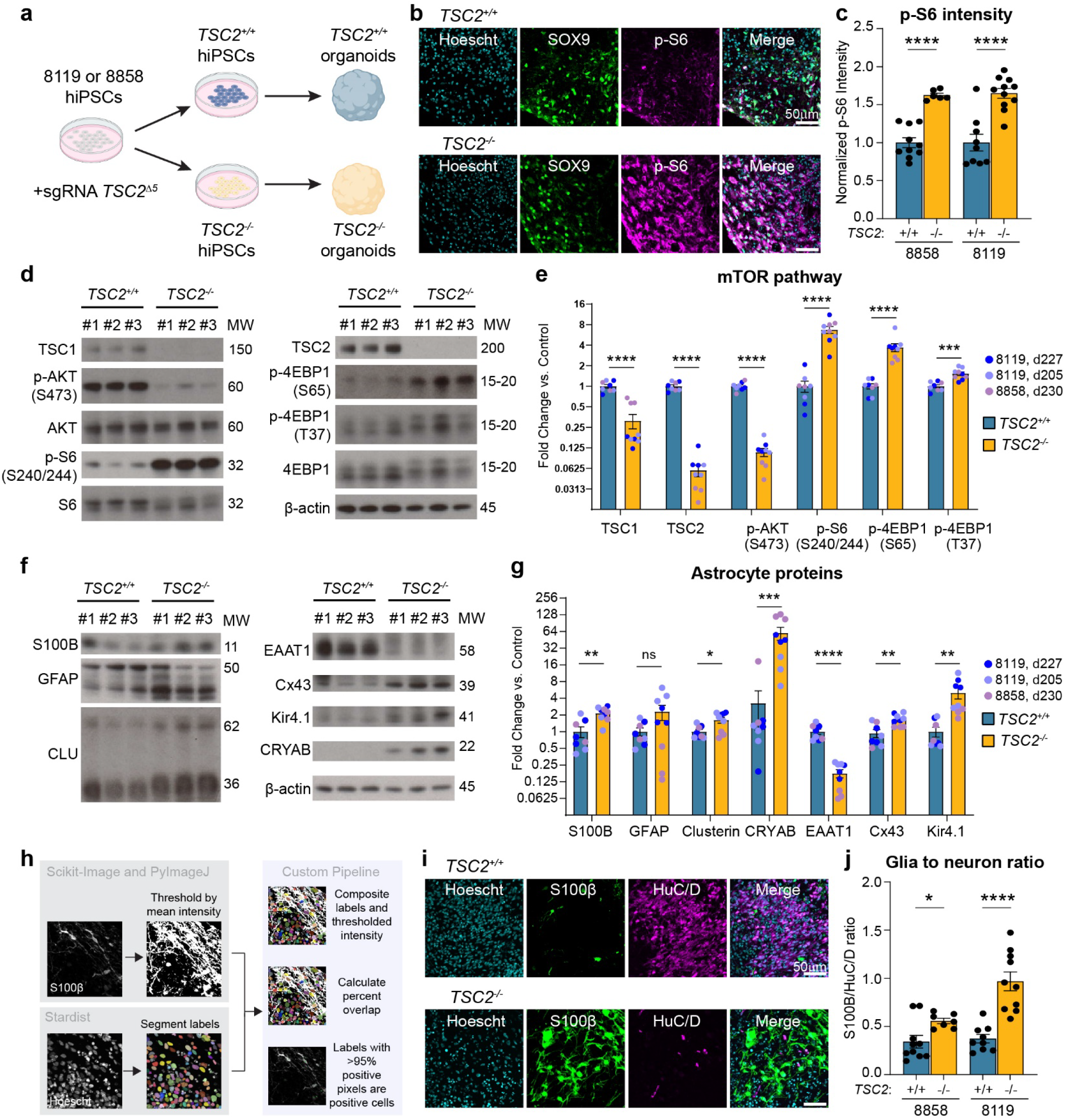
Protein expression changes in *TSC2^-/-^* brain organoids. **a,** Schematic of the gene editing and brain organoid differentiation approach. **b,** Example images of SOX9 and p-S6 immunostaining in *TSC2^+/+^* (top) and *TSC2^-/-^* (bottom) organoids at day 132 (8119 line). **c,** Quantification (mean ± SEM) of p-S6 intensity in cells from *TSC2^+/+^* and *TSC2^-/-^* organoids. Dots represent values for individual organoids. **d,** Example western blots of mTOR pathway proteins, showing three independent samples per genotype. MW denotes the approximate molecular weight. **e,** Quantification (mean ± SEM) of western blot data. Dots represent values for individual organoids. For statistics, sample sizes, and batches see Supplemental Table 10. **f,** Example western blots of astrocyte-related proteins. **g,** Quantification (mean ± SEM) of western blot data. Dots represent values for individual organoids. **h,** Schematic of the binary thresholding method used to determine if a cell is positive for a given marker. **i,** Example images of S100β and HuC/D immunostaining in *TSC2^+/+^* (top) and *TSC2^-/-^* (bottom) organoids at day 132 (8119 line). **j,** Quantification (mean ± SEM) of the glia to neuron ratio, as determined by the ratio of S100β and HuC/D positive cells in *TSC2^+/+^* and *TSC2^-/-^* organoids. Dots represent individual organoids.

To assess the morphology and protein expression profile of astrocytes specifically, we performed immunopanning from day ∼210-350 *TSC2^+/+^* and *TSC2^-/-^* organoids, which generated highly pure 2D astrocyte cultures (Fig. 4a,b). Western blotting of immunopanned astrocytes revealed robust protein-level changes in S100B, GFAP, clusterin, and CRYAB (Fig. 4c,d). We also observed altered expression of key astrocytic proteins including the excitatory amino acid transporter 1 (EAAT1 or GLAST), the gap junction protein connexin 43, and the potassium channel Kir4.1 (Fig. 4c,d). Alterations in these proteins can affect the functional properties of astrocytes, affecting their ability to uptake glutamate, mediate intercellular communication, and buffer potassium^46^.

**Figure 4:**
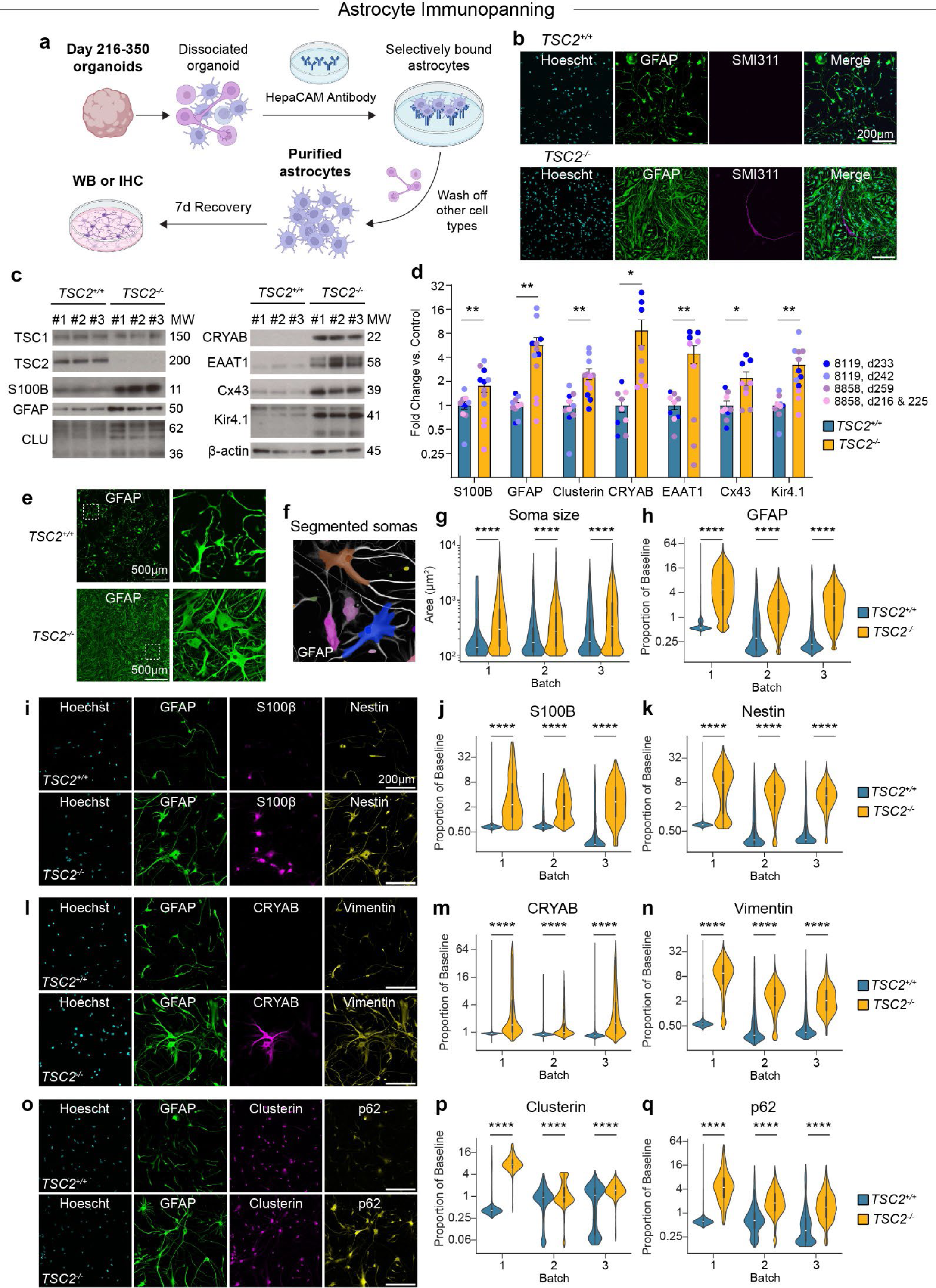
Purified astrocytes from *TSC2^-/-^* organoids have altered morphology and expression profiles. **a,** Schematic of the immunopanning process to isolate astrocytes from organoids. **b,** Example images of GFAP and SMI311 immunostaining in *TSC2^+/+^* and *TSC2^-/-^* immunopanned astrocytes from day 240 organoids from the 8119 hiPSC line. **c,** Example western blots of candidate DE proteins in immunopanned astrocytes, showing three independent samples per genotype. MW denotes the approximate molecular weight. **d,** Quantification (mean ± SEM) of western blot data, each dot represents one organoid. **e,** Example images of GFAP immunostaining showing altered morphology of *TSC2^-/-^* astrocytes immunopanned from day 240 organoids from the 8119 hiPSC line. **f,** Example images of segmented somas (translucent) and corresponding seed nuclei (solid). **g,** Violin plots show quantification of soma size for *TSC2^+/+^* and *TSC2^-/-^* astrocytes. Violins show overall distributions, and the overlaid box-and-whisker plots show quartiles (dark lines) and the median (white lines). **h-k,** Example images (**i**) and violin plots showing quantification of GFAP (**h**), S100β (**j**), and Nestin (**k**) levels in *TSC2^+/+^* and *TSC2^-/-^* astrocytes. **l-n,** Example images (**l**) and violin plots showing quantifications of CRYAB (**m**), and Vimentin (**n**) in *TSC2^+/+^* and *TSC2^-/-^* astrocytes. **o-q,** Example images (**o**) and quantifications of Clusterin (**p**), and p62 (**q**) in *TSC2^+/+^* and *TSC2^-/-^* astrocytes. Experiments in panels f-q were performed on astrocytes purified from day 350 organoids from the 8119 hiPSC line. See Supplemental Table 10 for statistics and sample sizes.

It was previously shown that it takes >300 days for astrocytes in brain organoids to exhibit features of mature astrocytes^45^. Therefore, to study how loss of *TSC2* affected the properties of mature astrocytes, we cultured *TSC2^+/+^* and *TSC2^-/-^* organoids for ∼350 days and performed immunopanning. Compared to controls, *TSC2^-/-^* astrocytes were highly enlarged and dysmorphic (Fig. 4e). We computationally segmented and quantified the size of individual astrocyte cell bodies from day 350 astrocytes (Fig. 4f and Extended Data Fig. 6a) and found that *TSC2^-/-^* astrocytes had dramatically larger soma sizes than *TSC2^+/+^* cells (Fig. 4g). These data are consistent with previous observations^26^, and reminiscent of dysmorphic cells observed in patient tubers. Related to this, we also observed multinucleated GFAP positive cells (from 2 up to 17 nuclei per cell) that were present in *TSC2^-/-^* cultures but only rarely seen in *TSC2^+/+^* cultures (Extended Data Fig. 6b). It is known that giant/balloon cells in tubers are often multinucleated^47,48^ and our data suggest that the multinucleated cells could be astrocytic in origin.

We performed immunostaining to assess the cellular expression of proteins encoded by candidate genes from our sequencing data. We found that *TSC2^-/-^* astrocytes had greater expression of reactive astrocyte proteins including GFAP, S100B, nestin, CRYAB, vimentin, and clusterin, at the level of individual cells (Fig. 4h-p). In line with the sequencing data, we also observed upregulation of p62 and APOE in *TSC2^-/-^* astrocytes (Fig. 4q and Extended Data Fig. 6c,d). These protein changes were reproducibly observed in independent astrocyte cultures obtained from day 240 *TSC2^+/+^* and *TSC2^-/-^* organoids (Extended Data Fig. 6g-r). Notably, immunopanned *TSC2^-/-^* astrocytes did not exhibit increased proliferation, as the percentage of Ki67+ cells was not different compared to controls (Extended Data Fig. 6e,f).

Together, these results show that across cell lines and organoid batches, many of the genes that were DE in the scRNAseq data were also increased at the protein level in purified astrocytes. The dramatic changes in astrocyte morphology and multinucleation are consistent with astrogliosis^49^, showing that loss of TSC2 drives cell autonomous reactive gliosis.

### Tuber cells display the reactive astrocyte signature found in *TSC2 -/-* cells *in vitro*

The analyses above identified a panel of DE genes and proteins in *TSC2^-/-^* cells in organoids. To test if these are also altered in tuber cells, we performed immunostaining on two tuber samples from individuals with TSC who had undergone resection surgery for seizure control (Supplemental Table 9). Compared to neighboring cells with low p-S6 levels, cells with high p-S6 in tubers had increased expression of four of the top DE proteins identified in our organoids: APOE (Fig. 5a,c), clusterin (Fig. 5b,d), CRYAB (Fig. 5e,g), and p62 (Fig. 5f,h), confirming the validity of the organoid model.

**Figure 5:**
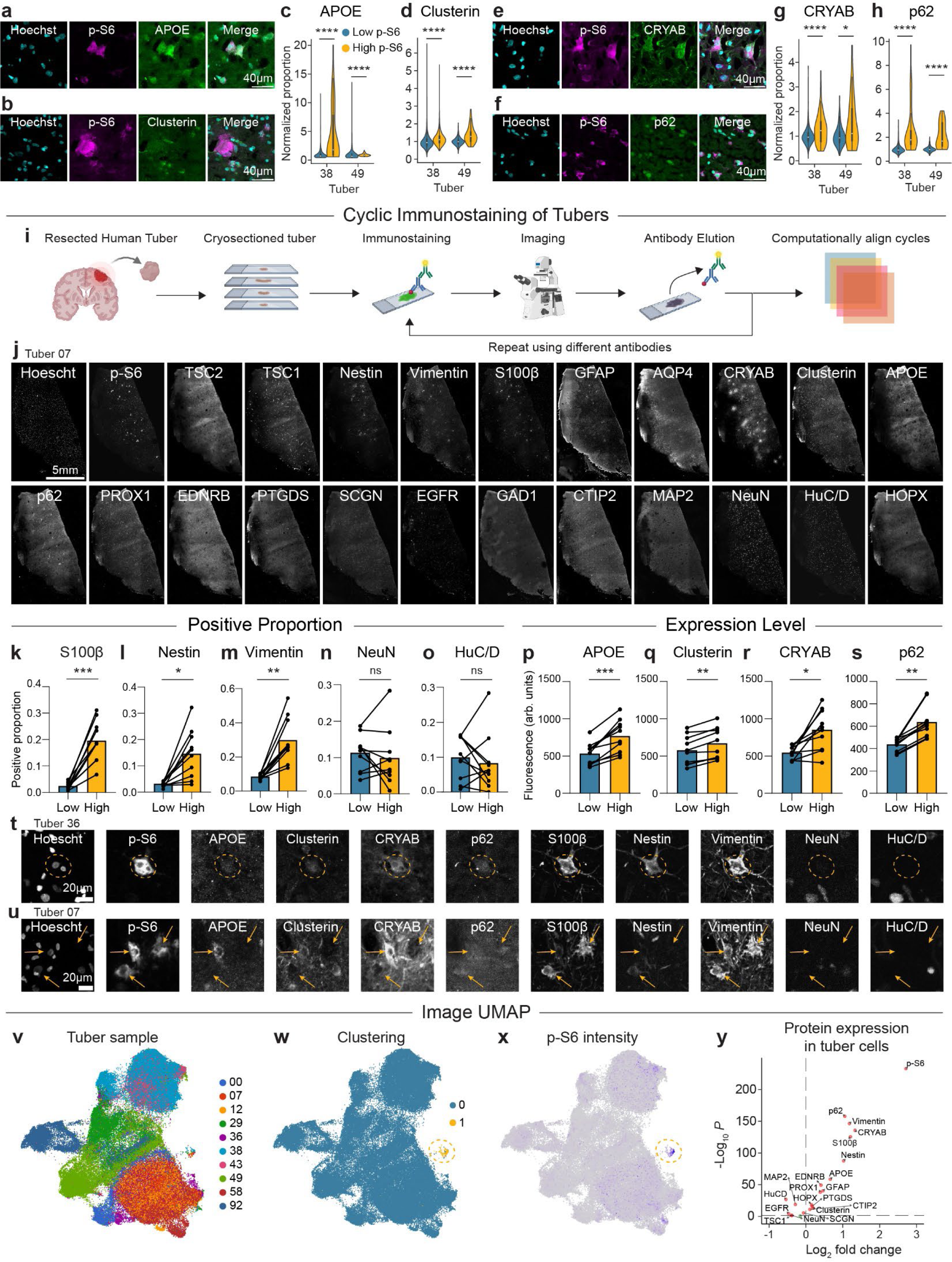
High p-S6 cells in TSC patient tubers express reactive astrocyte proteins. **a-h,** Example images and quantifications of p-S6 levels and candidate DE proteins per cell: APOE (**a,c**), Clusterin (**b,d**), CRYAB (**e,g**), and p62 (**f,h**). Violin plots show overall distributions, and the overlaid box-and-whisker plots show quartiles (dark lines) and the median (white lines). **i,** Schematic of the 4i cyclic staining protocol. **j,** Example images of a single tuber section (tuber ID #07) stained using 23 different antibodies. **k-o,** Graphs showing the proportion of cells positive for different proteins between cells with high p-S6 (“high”) and all other cells (“low”) within the same tuber for S100β (**k**), Nestin (**l**), Vimentin (**m**), NeuN (**n**), and HuC/D (**o**). Bars represent the mean. **p-s,** Intensity comparisons between cells with high p-S6 and all other cells within the same tuber for APOE (**p**), Clusterin (**q**), CRYAB (**r**), and p62 (**s**). n=10 tubers per comparison. Reported p-values are adjusted using the Bonferroni correction. **t,u** Example images of tuber cells across selected channels from two different TSC patients. **v,w**, UMAP derived from individual cell images across all channels and tubers, plotted by tuber sample (**v**) and by unbiased clustering (**w**). **x**, Feature plot of p-S6 intensity. **y**, Volcano plot of differential intensity between cells in cluster 1 (high p-S6) and cluster 0 (all others). Proteins more highly expressed in cluster 1 cells have a positive Log_2_ fold change, and proteins more highly expressed in cluster 0 cells have a negative Log_2_ fold change. See Supplemental Table 10 for statistics and sample sizes.

To gain a more comprehensive understanding of protein expression changes in tuber cells, we used 4i^50,51^, a multiplexed cyclic immunostaining approach, to probe the same slice of tuber tissue with dozens of antibodies (Fig. 5i,j) and computationally align the same cells across each cycle^52,53^ (Extended Data Fig. 7). In our antibody panel, we included our top DE candidates, general neuron and glia markers, and proteins identified in tubers from a recent study^25^. Cyclic staining of 10 tubers from 8 different TSC patients (Supplemental Table 9) revealed that cells with high p-S6 were more likely to be astrocytes and express candidate DE proteins compared to low p-S6 cells in the same tuber. Specifically, high p-S6 cells were more likely to be positive for the astrocytic proteins S100B, nestin, and vimentin, but were not more likely to be positive for the neuronal markers NeuN or HuC/D (Fig. 5k-o). Accordingly, high p-S6 cells had higher expression levels of APOE, clusterin, CRYAB, and p62 (Fig. 5p-s). We note that while these broad trends were found across tubers, individual tuber cells displayed varied protein expression profiles, even between adjacent cells (Fig. 5t,u and Extended Data Fig. 8a-e). Notably, in certain samples, we observed that cells with high p-S6 were associated with regions of lower TSC2 signal intensity (Extended Data Fig. 8f,g), suggesting cell autonomous loss of TSC2 protein.

The analyses above were based on comparisons of high versus low p-S6 cells in tubers. To examine protein expression trends across tuber cells in an unbiased manner, we developed an “image UMAP” approach. We established a workflow to extract the intensities and pixel relationships from 117,901 cells across all tubers and performed UMAP dimensionality reduction on this data (Fig. 5v and Extended Data Fig. 7a). We found that while different tubers generally separated in UMAP space, unbiased clustering identified a unified outlying cluster of 385 cells from 9 of the 10 tubers (Fig. 5w). When mapped by p-S6 intensity, this small cluster contained the cells with the highest p-S6 levels (Fig. 5x).

Differential intensity analysis between the cells in the outlying cluster versus all other cells showed that their expression profile aligned with the reactive astrocyte profile identified in *TSC2^-/-^* cells *in vitro*. These tuber cells had higher expression of p62, vimentin, CRYAB, S100B, nestin, APOE, GFAP, PTGDS and clusterin (Fig. 5y). Additionally, cells in the high p-S6 cluster had reduced expression of neuronal proteins including HuC/D, MAP2, and NeuN (Fig. 5y).

In summary, we find an abundance of reactive astrocytes and a paucity of neuronal markers in the hyperactive mTORC1 cells from TSC patient tubers using multiple methods for assessing protein expression. Our novel UMAP approach identifies a small, unique population of abnormal astrocytic cells within 9 out of 10 tubers that have hyperactive mTORC1 signaling. Taken together with our data in organoids, this suggests a model whereby cells with activated mTORC1 signaling are driven to become astrocytes, which are dysmorphic and express proteins that are indicative of a reactive state.

## Discussion

In this study, we show that *TSC2^-/-^* cells in human brain organoids preferentially generate astrocyte-lineage cells with transcriptional and morphological hallmarks of disease-associated reactivity. These astrocytes are morphologically similar to balloon cells in tubers, with dramatically larger somas and occasional multinucleation, indicating that balloon cells may be astrocytic in origin. In tubers surgically resected from individuals with TSC, we found that the small subset of cells with high mTORC1 signaling had a protein expression profile similar to *TSC2^-/-^* cells in organoids. These results indicate that reactive astrocytes can arise early in development due to mTORC1 hyperactivity and are positioned to be significant drivers of pathophysiology in TSC.

Glial pathology has been consistently reported in tubers^11,46^, but it has been unclear whether this was a consequence or a cause of chronic seizures and epilepsy. Our results in brain organoids show that activation of mTORC1 through biallelic loss of *TSC2* is sufficient to increase astrocyte differentiation and induce reactivity at the earliest stages of gliogenesis, in conditions where surrounding cells are not reactive. Notably, in organoids, this occurs in the absence of microglia, blood, or immune cells, which can induce astrogliosis in response to injury or disease^27^. These findings are congruent with recent studies showing that pro-gliogenic factors converge on mTORC1 signaling during human astrocyte development^54^ and that cytokine-induced astrocyte reactivity triggers mTORC1 activation and mTOR-dependent remodeling of the endolysosomal system in diseased astrocytes^55^. Together with our work, these studies provide a compelling case for the idea that mTORC1 activation is both pro-gliogenic and promotes the transition to a reactive state in response to disease-associated insults. In TSC, mTORC1 is autonomously activated due to loss of *TSC1/2* and therefore drives both astrocyte production and reactivity early in development, with likely consequences for neuronal development and function.

Our proposed model is consistent with other observations across the TSC and epilepsy literature. While much work in TSC has focused on neurons, which are also impacted^1,56^, abnormal glial cells and balloon cells are abundant in tubers^57^. Several studies in mouse models have identified glial abnormalities and astrocyte dysfunction resulting directly from Tsc1/2 loss in these cells^46,58^. Reactive astrocytes could cause seizures through failure of homeostatic functions, including glutamate uptake and potassium buffering, or they could gain detrimental functions that cause blood brain barrier disruption or trigger secondary neuroinflammation^59^. In other contexts, such as acquired epilepsy, the presence of reactive astrocytes alone is sufficient to drive epileptogenesis^59^. If the mTORC1-activated cells in tubers are indeed reactive astrocytes, this model would explain how a small population of cells is able to generate a localized tuber while also affecting cortical networks more broadly. It would also explain why non-imaging-based methodologies, such as bulk sequencing, may fail to detect second-hit mutations, as in our analysis, the mTORC1 activated cells only represent 0.3% of the overall population and can be easily missed. It is also possible that the “second-hit” may not be a canonical LOH mechanism but could be any stochastic genetic or epigenetic event during neural development that drives mTORC1 activation over a critical threshold.

Finally, our work hints at common underlying factors connecting developmental disorders, like TSC, and neurodegenerative diseases such as Alzheimer’s disease. Genetic variants of many of the DE genes identified in this study, such as *CLU* and *APOE,* have been associated with increased Alzheimer’s disease risk^28,29^. Moreover, mTORC1 dysregulation has been reported in Alzheimer’s disease^1^. Alterations in *SQSTM1* expression and autophagy have also been implicated broadly in neurodegenerative disorders^60^. There is a growing body of work demonstrating the role of reactive glia, both astrocytes and microglia, in driving pathophysiology in these diseases^61^. Activation of mTORC1 signaling, either cell autonomously due to loss of TSC1/2 complex function, or by some other means, may drive astrocytes to an aberrant state that contributes to multiple neurological disorders. Future study of the mechanisms by which mTORC1 contributes to astrocyte reactivity, as well as how this state may be reversed, will lead to deeper insights about the role of glia in establishing and maintaining brain function, with important therapeutic potential.

## Author Contributions

T.L.L. and J.D.B. generated cell lines and organoids and performed single-cell RNA sequencing experiments. T.L.L. conducted code-based analyses. T.Y. performed immunopanning and Western blotting experiments. G.A.G performed the surgical resection of tuber samples, which were collected and shared by and B.E.P. D.H. and H.S.B. supervised the project.

## Acknowledgements

This work was supported by R01NS097823 and a Siebel Stem Cell Center Seed Grant (to H.S.B.). T.L.L. was supported by a postdoctoral fellowship from CIRM Training Program EDUC4-12790. J.D.B. was supported by a Predoctoral Award from the American Epilepsy Society and a Frederick Banting and Charles Best Canada Graduate Scholarship from the Canadian Institutes for Health Research (#356733). D.H. was supported by a Chan Zuckerberg Biohub Investigator award and the Siebel Stem Cell Center. H.S.B. was supported by a Chan Zuckerberg Biohub Investigator award. Sequencing experiments were performed by the Chan Zuckerberg Biohub San Francisco Genomics Platform led by Dr. Norma Neff and QB3 Genomics, UC Berkeley, Berkeley, CA RRID:SCR_022170.

**The authors declare no competing interests.**

## Extended Data Figures

**Extended Data Fig. 1:**
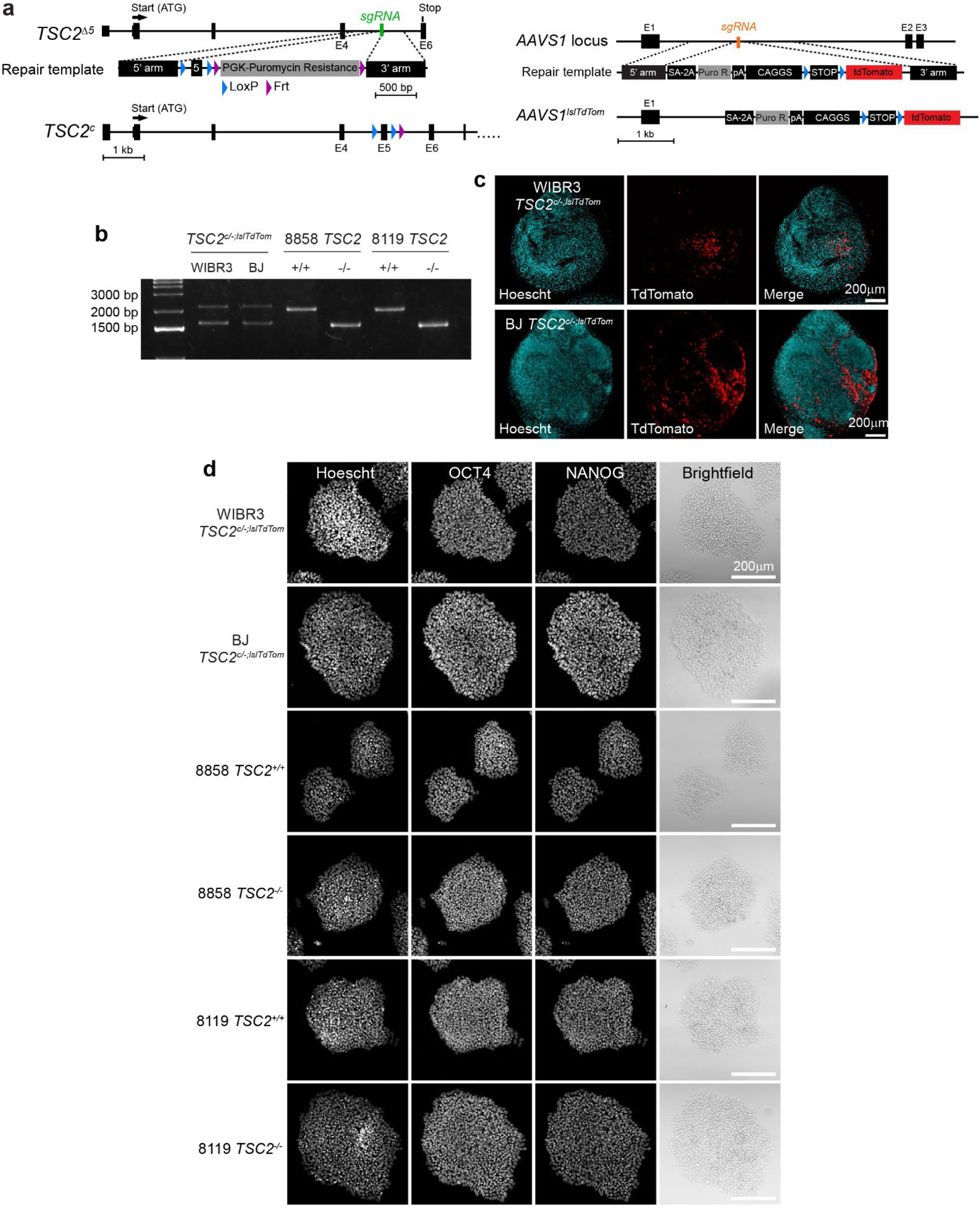
Validation of mosaic organoids, stem cell pluripotency, and gene editing. **a,** Schematic of the *TSC2^c/-;LSL-TdTom^* genetic system, showing the insertion of the floxed exon 5 to generate a conditional *TSC2* allele (left), together with a Cre-dependent TdTomato fluorescent reporter in the *AAVS1* safe harbor locus (right). **b,** PCR genotyping for the *TSC2* exon 5 deletion. The WT allele produces a 2000 bp product, the floxed exon 5 allele produces a 2070 bp product, and the exon 5 deleted allele produces a 1500 bp product. **c,** Example images of immunostained organoid sections showing TdTomato-positive cells in WIBR3 and BJ *TSC2^c/-;LSL-TdTom^* organoids at day 79 (WIBR3) and day 85 (BJ). **d,** Images of human pluripotent stem cell colonies immunostained for the pluripotency markers OCT4 and NANOG in feeder-free cultures.

**Extended Data Fig. 2:**
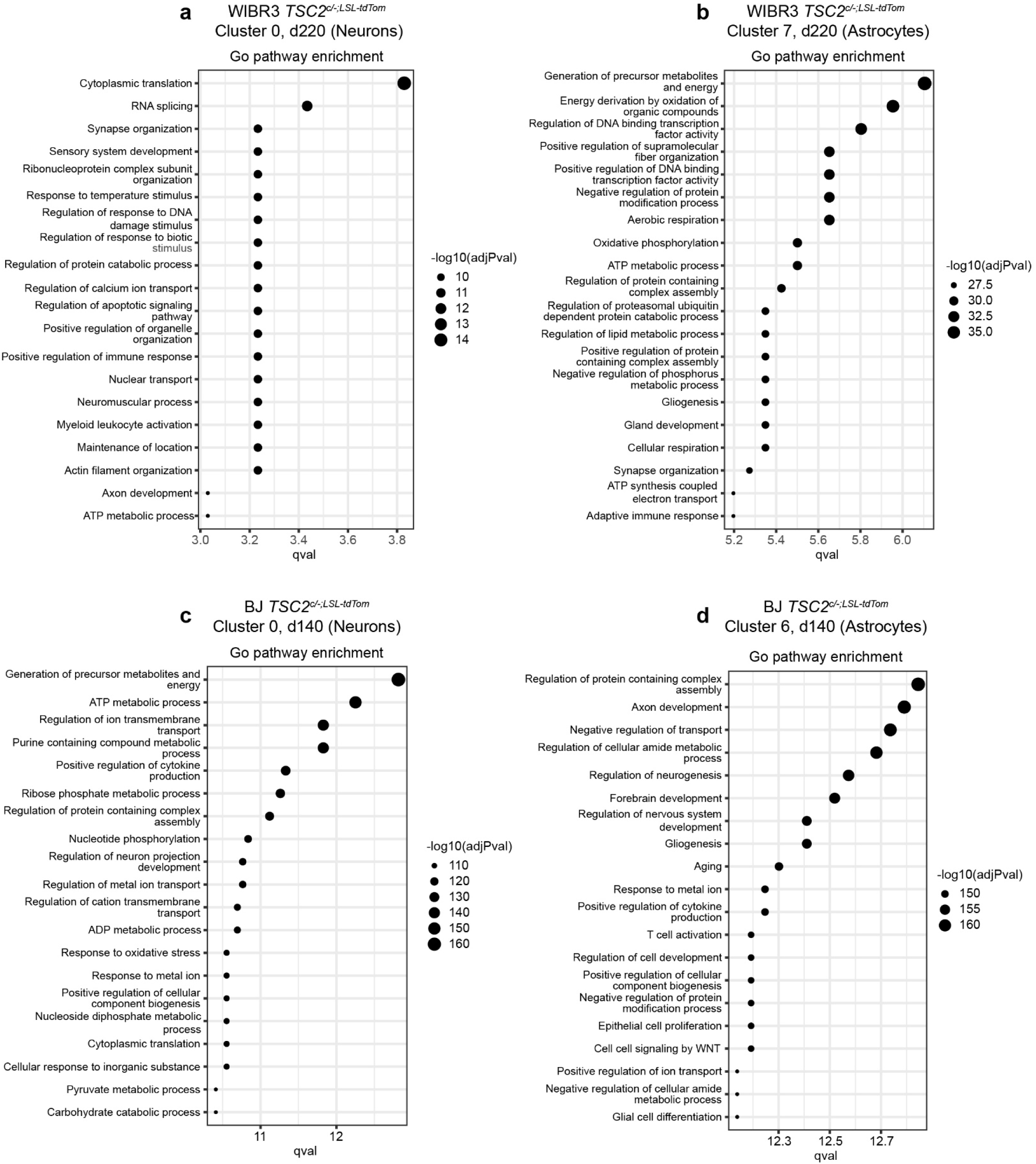
Pathway analysis of single-cell RNA sequencing data. **a-d:** Enriched pathways derived from Single Cell Pathway Analysis for the largest neuron cluster **(a)** and an astrocyte cluster **(b)** from WIBR3 *TSC2^c/-;LSL-TdTom^* organoids, and the largest neuron cluster **(c)** and an astrocyte cluster **(d)** from BJ *TSC2^c/-;LSL-TdTom^* organoids.

**Extended Data Fig. 3:**
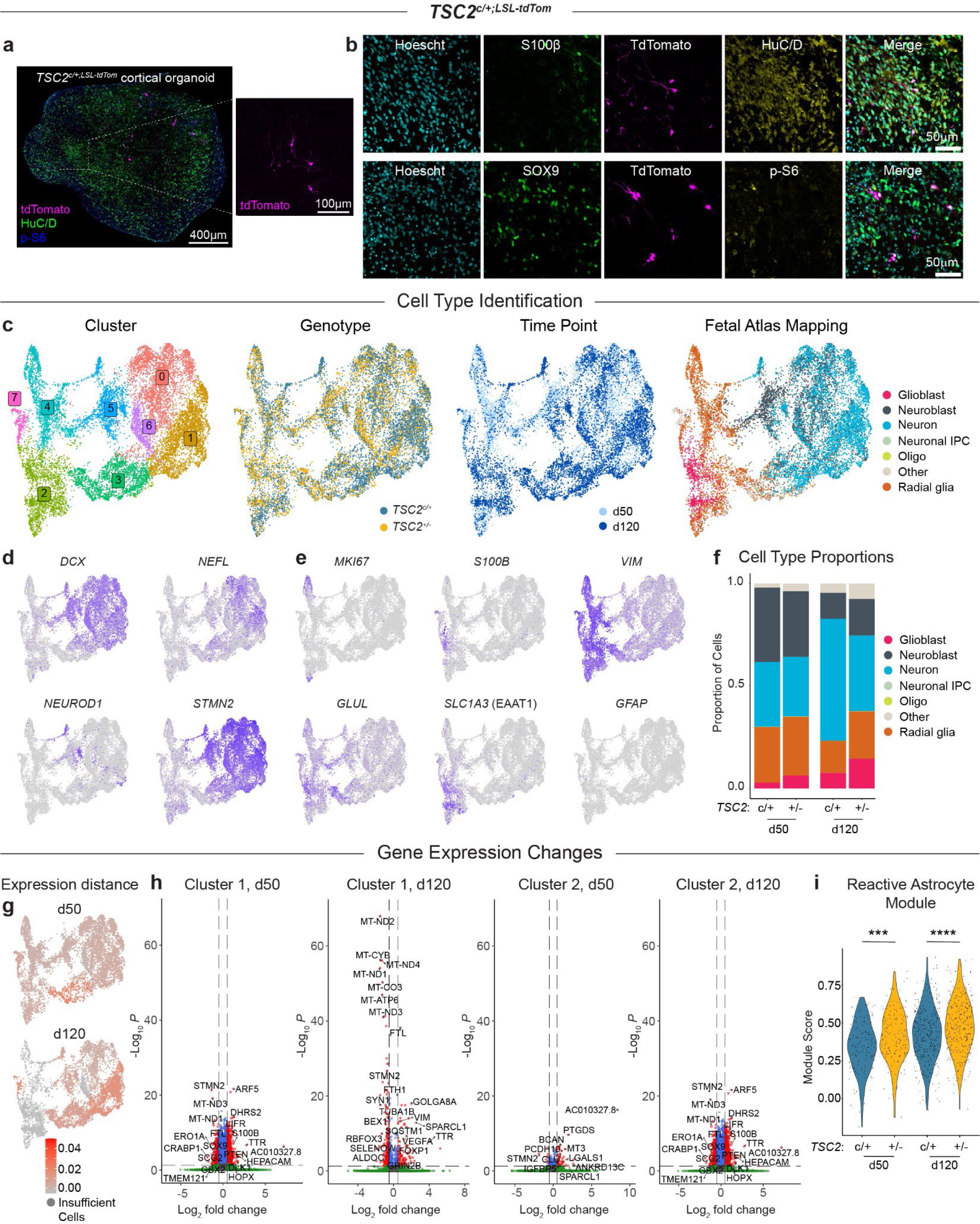
Single-cell RNA sequencing of WIBR3 *TSC2^c/+;LSL-TdTom^* brain organoids. **a,** Example image of an immunostained WIBR3 *TSC2^c/+;LSL-TdTom^* organoid showing tdTomato-positive *TSC2^+/-^* cells. HuC/D labels neurons in green and p-S6 is a read-out for mTORC1 activity in blue. **b,** Example images of day 120 *TSC2^c/+;LSL-TdTom^* organoid sections immunostained for S100β, TdTomato, and HuC/D (top images) or SOX9, TdTomato, and p-S6 (bottom images). **c,** Brain organoids were generated from *TSC2^c/+;LSL-TdTom^* hESCs and exposed to Cre lentivirus at day 8. Organoids were cultured until day 50 or 120, at which time they were dissociated, separated by FACS, and processed for 10x scRNA-seq. UMAP plots of scRNA-seq results, showing cells grouped by unbiased clustering, genotype, time point, and mapping to a fetal brain atlas. **d,** Feature plots of selected neuronal genes. **e,** Feature plots of selected astrocyte and progenitor genes. **f**, Cell type proportions, as assayed by atlas mapping, divided by genotype and time point. **g**, Cluster-based normalized expression distances between *TSC2^c/+^* and *TSC2^+/-^* cells. Expression distances were calculated between the two genotypes within each cluster. **h,** Volcano plots of differential gene expression between *TSC2^c/+^* and *TSC2^+/-^* cells in the astrocyte cluster (cluster 1) and a neuronal cluster (cluster 2) at day 50 and day 120. Genes more highly expressed in *TSC2^+/-^* cells have a positive Log2 fold change, and genes more highly expressed in *TSC2^c/+^* cells have a negative Log2 fold change. **h,** Violin plots show module scores for a reactive astrocyte gene module divided by genotype. Violins show overall distributions and dots represent values for individual cells. See Supplemental Table 10 for statistics and sample sizes.

**Extended Data Fig. 4:**
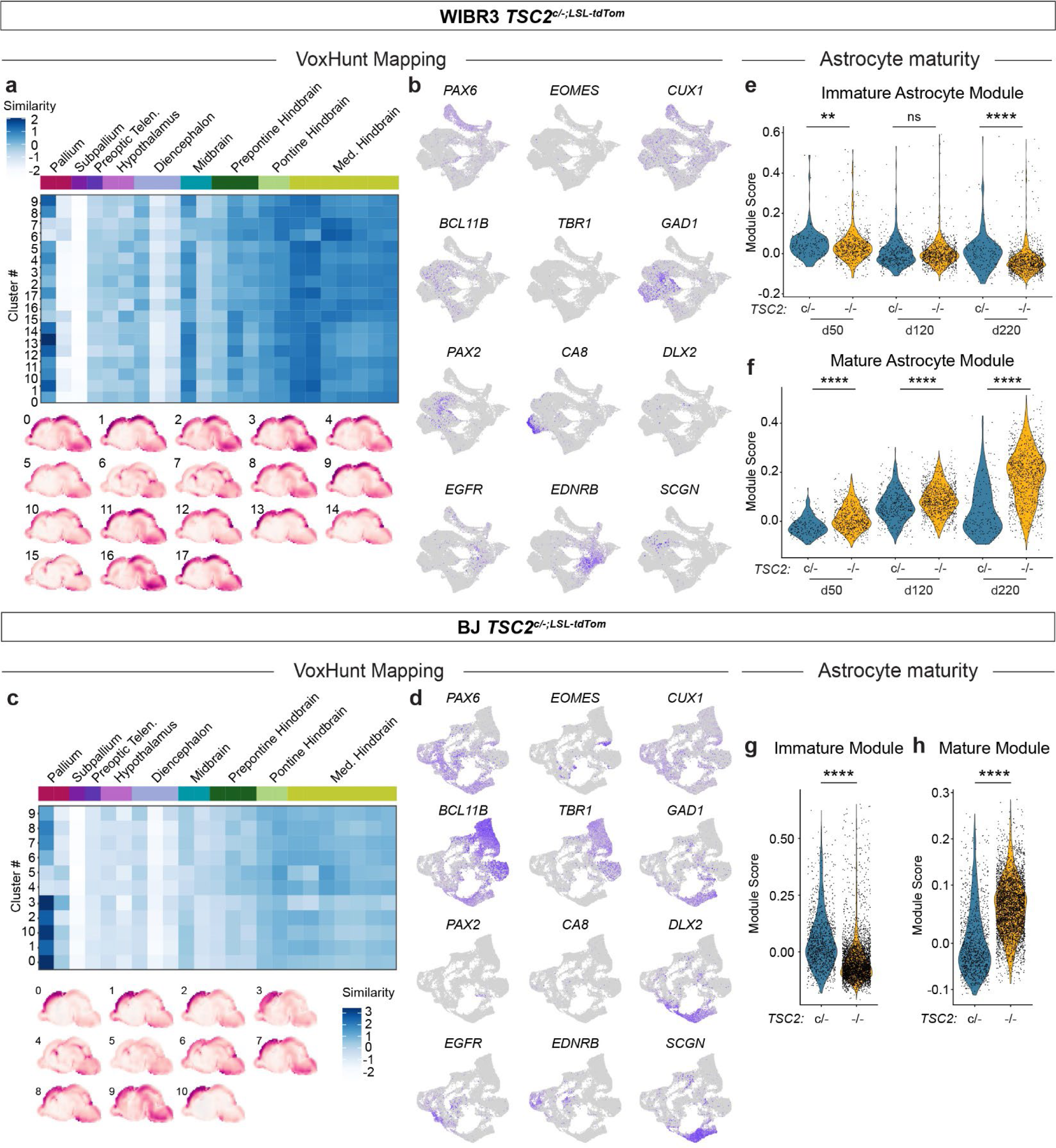
Additional single-cell RNA sequencing analyses. **a,** Spatial similarity mapping of each cluster for the WIBR3 *TSC2^c/-;LSL-TdTom^* scRNA-seq dataset to the E18.5 mouse brain via VoxHunt. Bottom, spatial similarity maps for each cluster projected onto E18.5 sagittal mouse brains. **b,** Feature plots of genes related to neuronal subtype and regional identity in WIBR3 *TSC2^c/-;LSL-TdTom^* organoids. **c,** Spatial similarity mapping of each cluster for the BJ *TSC2^c/-;LSL-TdTom^* scRNA-seq dataset to the E18.5 mouse brain via VoxHunt. Bottom, spatial similarity maps for each cluster projected onto E18.5 sagittal mouse brains. **d,** Feature plots of genes related to neuronal subtype and regional identity in BJ *TSC2^c/-;LSL-TdTom^* organoids. **e,f,** Violin plots show module scores calculated for immature **(e)** and mature **(f)** astrocyte gene modules in WIBR3 *TSC2^c/-;LSL-TdTom^* organoids, divided by time point and genotype. Violins show overall distributions and dots represent values for individual cells. **g,h,** Module scores calculated for immature **(g)** and mature **(h)** astrocyte gene modules in BJ *TSC2^c/-;LSL-TdTom^* organoids, divided by genotype. See Supplemental Table 10 for statistics and sample sizes.

**Extended Data Fig. 5:**
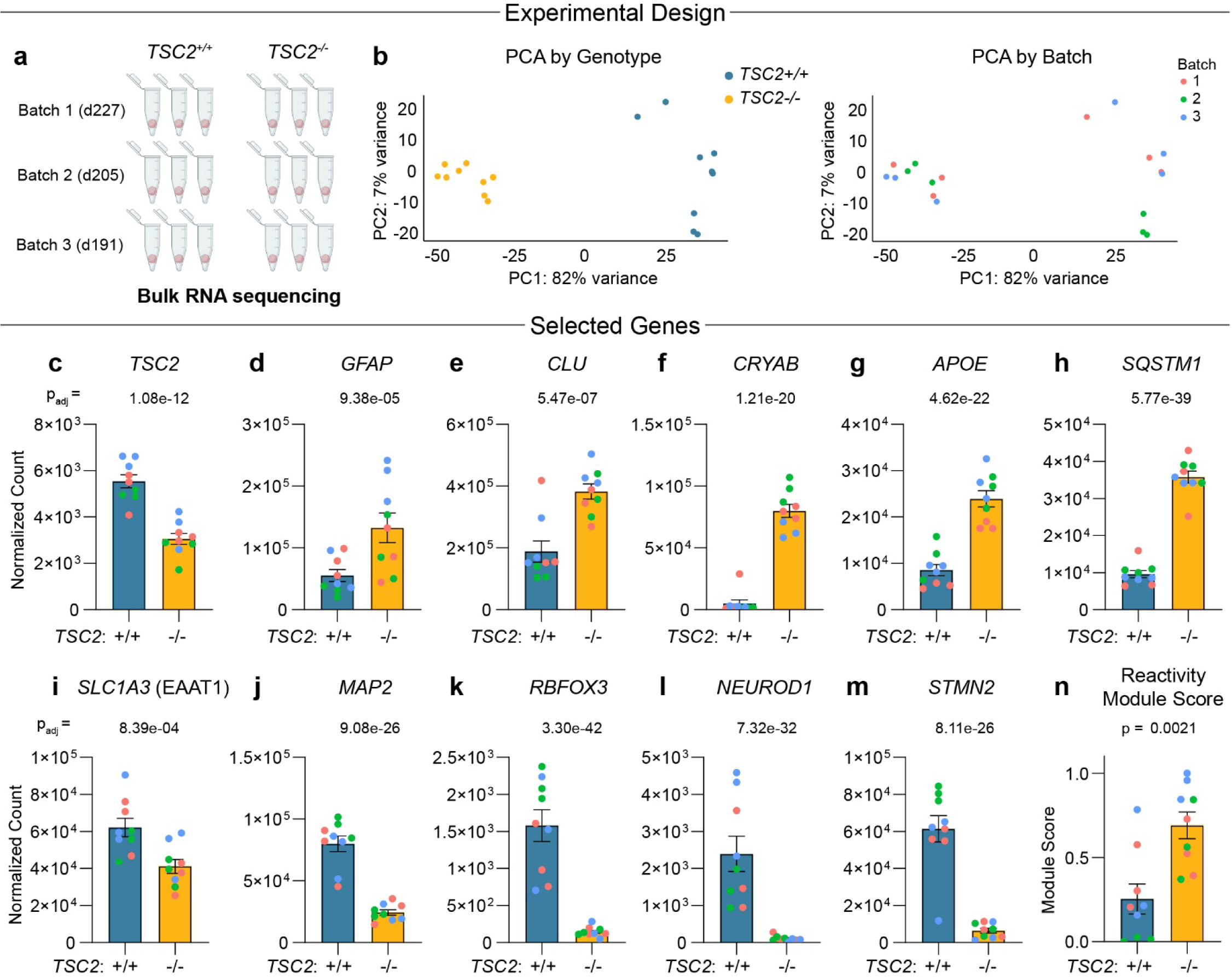
Bulk RNA sequencing of *TSC2^+/+^* and *TSC2^-/-^* organoids. **a,** Experimental schematic of organoid batches and genotypes collected for bulk RNA sequencing (differentiated from the 8119 hiPSC line). **b,** PCA plot of gene expression across all 18 samples, colored by genotype (left) or batch (right). **c-m,** Quantification (mean +/- SEM) of gene expression changes for selected candidate genes. Adjusted P values were calculated using the DESeq2 differential expression analysis. **n,** Quantification (mean +/- SEM) of the reactive astrocyte gene module score calculated for each sample. The P value was calculated using the Wilcoxon rank-sum test. For panels **c-n**, dots represent values for individual organoids.

**Extended Data Fig. 6:**
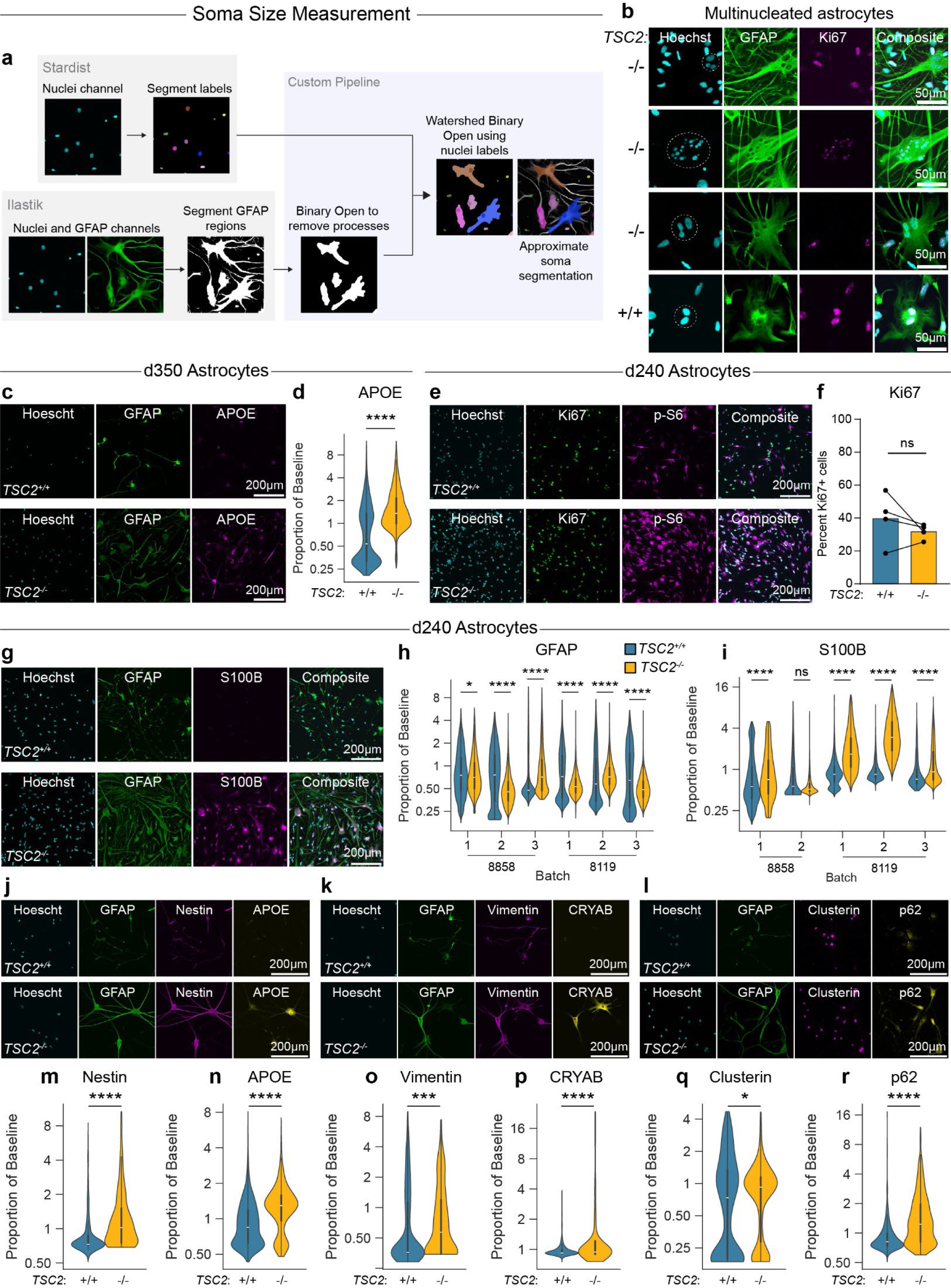
Additional analysis of immunopanned astrocytes. **a,** Schematic of the soma size analysis pipeline, including segmentation of the nuclei and GFAP channels. **b,** Selected multinucleated cells from *TSC2^-/-^* and *TSC2^+/+^* immunopanned astrocytes, immunostained for GFAP and the proliferation marker Ki67. **c,** Example images of GFAP and APOE immunostaining in astrocytes purified from day 350 *TSC2^+/+^* and *TSC2^-/-^* organoids (8119 hiPSC line). **f,** Violin plots show quantification of APOE levels per cell. Violins show overall distributions, and the overlaid box-and-whisker plots show quartiles (dark lines) and the median (white lines). **e,** Example images of Ki67 and p-S6 immunostaining of astrocytes purified from day 240 *TSC2^+/+^* and *TSC2^-/-^* organoids. **f,** Quantification of the proportion of Ki67-positive cells in *TSC2^+/+^* and *TSC2^-/-^* astrocyte cultures isolated in parallel from day 240 organoids. Bars represent the mean and dots represent values for individual batches. A total of 66,308 cells were analyzed across 8 cultures (4 cultures per genotype). **g,** Example images of GFAP and S100β immunostaining in astrocytes purified from day 240 *TSC2^+/+^* and *TSC2^-/-^* organoids (8119 hiPSC line). **h,i** Violin plots show quantification of GFAP (**h**) and S100β (**i**) levels per cell across batches and hiPSC lines. **j-l** Example immunostaining images of candidate DE proteins in astrocytes purified from day 240 *TSC2^+/+^* and *TSC2^-/-^* organoids (8858 and 8119 hiPSC lines). **m-r**, Violin plots show quantification of Nestin (**m**), APOE (**n**), Vimentin (**o**), CRYAB (**p**), Clusterin (**q**), and p62 (**r**) levels per cell. For sample sizes, P values, and statistical tests of intensity-based measurements, see Supplemental Table 10.

**Extended Data Fig. 7:**
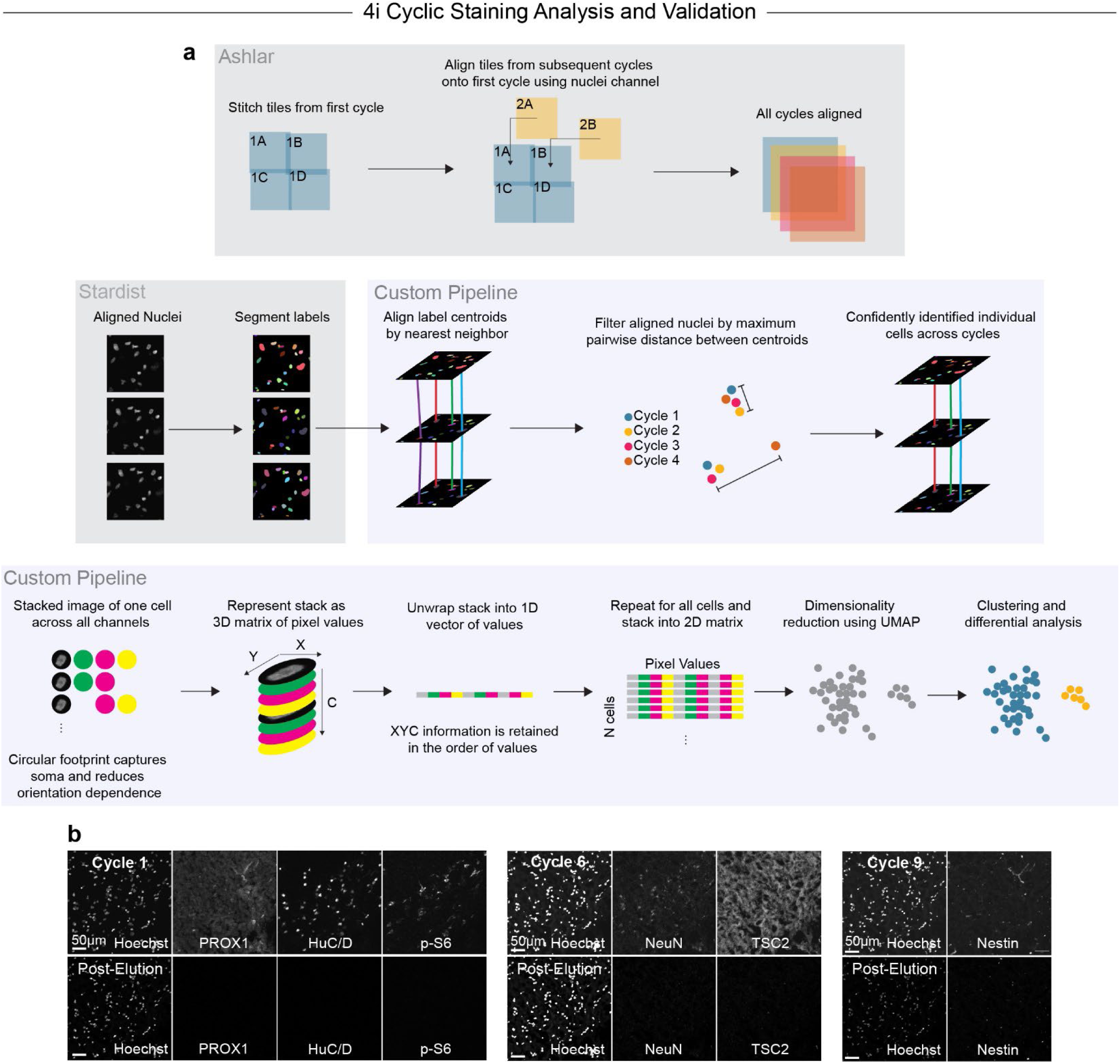
Cyclic staining analysis pipeline and validation. **a,** Schematic of the analysis pipeline for aligning cells across cycles and generating the image UMAP. Tiles from the first cycle were stitched, then tiles from subsequent cycles were aligned to the first cycle using ASHLAR. Stardist was used to segment the nuclei channel of each cycle individually. For each segmented nucleus in the first cycle, the corresponding nucleus in subsequent cycles was found using nearest-neighbor analysis. After all nuclei were aligned, poor alignments were excluded using a threshold for the maximum distance between any two centroids across cycles for each nucleus. To generate the image UMAP, a 30-pixel radius around each centroid was selected for each cell and converted to a 1-D vector of values. These vectors were stacked to generate a 2-D matrix. This matrix was used to generate a UMAP projection for clustering and differential analysis. **b,** Example images of antibody elution across immunostaining cycles. After elution, samples were stained with Hoechst and re-imaged using the same exposure settings as the previous cycle to confirm that no residual signal remained.

**Extended Data Fig. 8:**
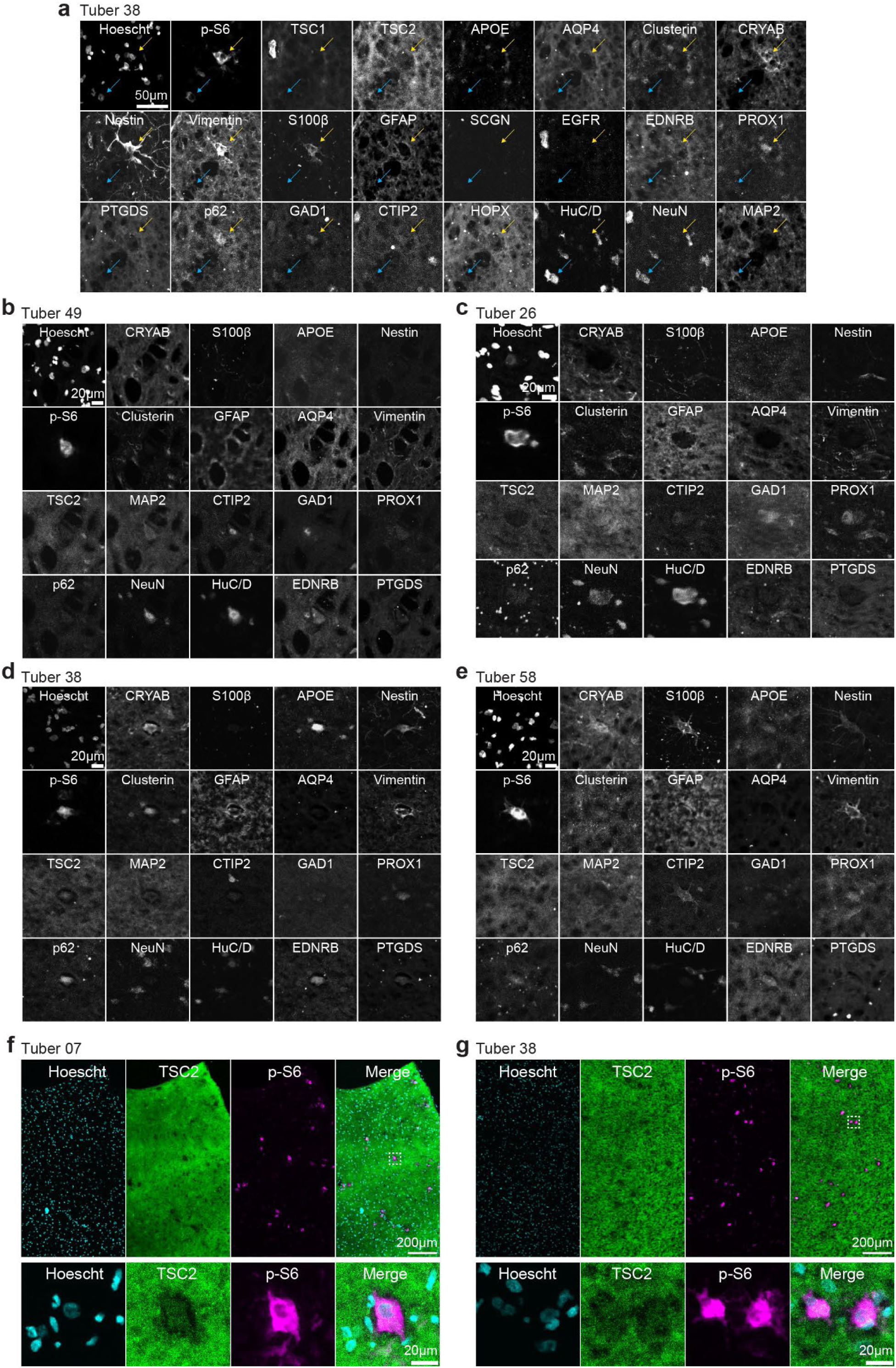
Additional cyclic staining example images from TSC patient tubers. **a,** Example images of a high p-S6 astrocyte (yellow arrow) and a high p-S6 neuron (blue arrow) across all antibodies. **b,c,** Two sets of example images of high p-S6 cells expressing neuronal markers from different tubers. **d,e,** Two sets of example images of multinucleated high p-S6 cells expressing astrocytic markers from different tubers. **f,g** Example images of two selected tubers immunostained for p-S6 and TSC2 at two scales, showing that the brightest p-S6 cells tend to have depletion of TSC2 immunoreactivity.

## Methods

### Human pluripotent stem cell (hPSC) culture

WIBR3 hESCs (NIH stem cell registry 0079) were initially obtained from Dr. Rudolf Jaenisch’s laboratory^1^. WIBR3 hESC lines were cultured according to published protocols^2,3^. hESCs were maintained on a layer of inactivated mouse embryonic fibroblasts (MEFs, CD-1 strain, Charles River) in hPSC medium, consisting of DMEM/F12 supplemented with 20% KnockOut Serum Replacement (Thermo Fisher, 10828010), 2 mM L-glutamine (Thermo Fisher, A2916801), 1% nonessential amino acids (Thermo Fisher 11140050), 0.1 mM 2-mercaptoethanol (Sigma, M6250), and 4 ng/ml fibroblast growth factor (FGF)-Basic (AA 1-155) recombinant human protein (Thermo Fisher, PHG0261). Cultures were passaged every 7 days with collagenase type IV (1.5 mg/ml; Thermo Fisher, 17104019) and gravitational sedimentation by washing 3 times in wash media composed of DMEM/F12 supplemented with 5% fetal bovine serum (FBS, Thermo Fisher, A5670801) and 1,000 U/ml penicillin/streptomycin (Thermo Fisher 15070063).

The BJ, 8858, and 8119 hiPSC lines were maintained in feeder-free conditions. iPSCs were cultured on TC-treated 6-well plates (Corning, 3516) coated with vitronectin (VTN-N, Gibco, A14700) and maintained in E8 media (Gibco, A1517001). Cultures were passaged using a 7-minute room temperature incubation with EDTA (Thermo Fisher, 15575020).

All cell lines were tested regularly for *Mycoplasma* contamination. On-target gene editing was confirmed by PCR (Extended Data Fig. 1b). Pluripotency status was confirmed by immunostaining with OCT4 and NANOG (Extended Data Fig. 1d). Genomic integrity of the stem cell lines was verified after gene editing using array comparative genomic hybridization (Cell Line Genetics) (Supplemental Table 11).

Gene editing of hESCs was performed using CRISPR/Cas9 editing and validated in our lab as previously reported^4^. Briefly, constitutive *TSC2* exon 5 deletion mutants were generated by electroporating cells with two px330 plasmids^5^ containing single-guide RNAs targeting the genomic regions of interest, as well as a GFP-encoding plasmid. After recovery, GFP-positive cells were selected using FACS, and single-cell-derived hESC colonies were manually picked, replated, and expanded.

WIBR3 *TSC2^c/-;LSL-TdTom^* hESCs, *TSC2^c/+;LSL-TdTom^* hESCs and BJ *TSC2^c/-;LSL-TdTom^* hiPSCs were generated using CRISPR/Cas9 gene-editing as previously described^4^. A single-guide RNA containing a *TSC2* exon 5 cassette flanked by loxP sites was cloned into px330, electroporated into *TSC2^+/-^* hESCs, and clonal colonies underwent puromycin selection. The puromycin resistance cassette was removed, and the Ai9 tdTomato Cre reporter cassette was added to the *AAVS1* safe harbor locus^6^ using the same genome editing methods described above.

All unique biological materials in this manuscript (e.g. gene edited human stem cell lines) will be provided to qualified users upon request and upon completion of the relevant material transfer agreements.

### Organoid differentiation

Cortical organoid generation was performed as described previously^7,8^. For feeder-based cultures, hESCs were isolated and removed from MEFs using accutase (Stemcell Technologies, 07920) for 20 minutes. The cell suspension was collected and strained through a 40 μm strainer in hESC media. This cell suspension was spun down for 5 minutes at 1000 rpm. The supernatant was removed and resuspended in 5 ml of hESC media without fibroblast growth factor 2 (FGF2), supplemented with 10 μm Y-27632 dihydrochloride (Selleckchem, S1049). Cells were counted and resuspended to a concentration of 2.7×10^6^ cells/ml and 6 ml of this suspension was deposited into one well of a 6-well Aggrewell 800 plate (Stemcell Technologies, 34825). After the aggregation of single cells into embryoid bodies (EBs) overnight, the EBs were removed and put into 10 cm ultra-low attachment dishes (Corning, 4615). On days 1–5, EBs were cultured in DMEM/F12 supplemented with 20% KnockOut Serum Replacement, 2 mM L-glutamine, 1% nonessential amino acids, 0.1 mM 2-mercaptoethanol, 1,000 U/mL penicillin-streptomycin, supplemented with 10 μM Dorsomorphin (Abcam, ab146597) and 10 μM SB-431542 (R&D Systems, 1614/10).

For feeder-free cultures, hiPSCs were cultured to high density (80-90% confluency). 24 hours before aggregation, hiPSCs were pretreated with 1% DMSO (Sigma, D2438) in E8 media. For aggregation, hiPSCs were isolated using a 7-minute incubation with accutase, and 3 million cells in 2 mL of E8 were transferred into one well of a 24-well Aggrewell 800 plate (Stemcell Technologies, 34815). The following day, aggregates were dislodged and transferred into 10 cm ultra-low attachment dishes, with aggregates from one well being transferred into two 10 cm dishes.

From day 1 to 5, organoids were cultured in E6 media (Thermo Fisher, A1516401), supplemented with 2.5 μM Dorsomorphin and 10 μM SB431542. After day 5, feeder-based and feeder-free cultures followed the same protocol. On day 6, organoids were cultured in neural induction media, consisting of Neurobasal-A (Thermo Fisher, 10888022), B-27 Supplement minus vitamin A (Thermo Fisher, 12587010), 50 U/mL penicillin-streptomycin, and 1X GlutaMAX (Thermo Fisher, 35050-061). During this time, neural induction media was supplemented with 20 ng/ml FGF (R&D, 236-EG) and 20 ng/ml EGF (R&D, 233-FB). A full media change was performed every day from days 6–15 and then every other day until day 25. From days 25–43, the organoids were grown in neural induction media supplemented with 20 ng/ml BDNF (Peprotech, 450-02) and 20 ng/ml NT-3 (Peprotech 450-03), with media changes every 4 days. From day 43 onward, organoids were maintained in neural induction media without BDNF or NT-3, with media changes every 4 days until harvest.

To generate the mosaic organoid models (*TSC2^c/-;LSL-TdTom^* and *TSC2^c/+;LSL-TdTom^*), on day 8 post-differentiation from hPSCs, organoids were transduced with UBC-Cre-RFP lentivirus (Kerafast, FCT224) by adding 5 uL of 1.0 x 10^8^ virus to each 10 cm dish containing approximately 20 organoids, with a media change after 24 hours.

### Organoid Dissociation for FACS

Dissociation of organoids for fluorescence-activated cell sorting (FACS) followed a protocol for dissociation of mouse cortex for primary neuronal culture^9^. First, dissociation media (DM) was made consisting of calcium and magnesium free HBSS (Invitrogen, 14185-052), 1 mM Sodium Pyruvate (Life Technologies, 11360070), 0.1% D-glucose (Sigma, G8769) and 10 mM pH 7.3 HEPES (Invitrogen, 15630-080). Next, the dissociation solution was made consisting of 5 ml DM, 172 ul Papain Solution (Worthington, LS003126), and 5.5mM L-Cysteine (Sigma, 168149). This solution was warmed at 37°C for 15 minutes and then filter-sterilized through a 0.22 μm filter. Organoids were transferred into DM and incubated at 37°C for 40 minutes. During this time, trypsin inhibitor (TI) solution was made consisting of 10 mg of Trypsin Inhibitor (Sigma) in 10 ml DM, pre-warmed at 37°C for >15 minutes and then filter-sterilized. After incubation, the intact organoid was washed twice with TI, then incubated in TI for 4 minutes at 37°C. During this time, the sorting buffer (SB) of 1x DPBS with calcium and magnesium (Thermo Fisher, 14040117) with 10 μM Y-27632 was made and placed on ice. After 4 minutes at 37°C, the TI was removed from the tube with the organoid and 2 ml of SB was added. The organoid was then mechanically dissociated by triturating 5-10 times through a 5 ml serological pipette within this solution. The dissociated cell solution was then taken up into the serological pipette and passed through a 70 μm cell strainer into a 50 ml conical tube. This passed-through solution was then placed into a polypropylene FACS tube on ice.

### FACS and single cell RNA sequencing

Dissociated cells were sorted on a BD Aria Fusion cell sorter with a 70 μm nozzle. When sorting for fluorophores, a negative control of a dissociated non-fluorophore labeled organoid was sorted first to ensure proper gating. After sorting, the cells were centrifuged at 300 x g for 5 minutes at 4 °C and then counted on a hemocytometer. Cells were then processed through the 10x Genomics 3’ single-cell sequencing pipeline for v2, v3, or v3.1 according to the manufacturer’s protocol. cDNA from the 10x protocol was assessed for quality at the UC Berkeley Functional Genomics Lab using an Agilent 2100 Bioanalyzer, and libraries were prepared using the 10x Genomics protocol. Sequencing was performed at the UC Berkeley Genomics Sequencing Laboratory (QB3 Genomics, UC Berkeley, Berkeley, CA, RRID:SCR_022170) or at the Chan Zuckerberg Biohub San Francisco Genomics Platform (Supplemental Table 12).

### Processing and analysis of single-cell sequencing data

FASTQ files were aligned to a modified version of the human genome (GRCh38) Cell Ranger 6.1.2 (10x Genomics). Cell Ranger gene expression matrix outputs were then loaded into Seurat 5.1^10^. Data from each individual sample was turned into a Seurat object and metadata regarding time point, genotype and batch was added to each object. Each object was subset, extracting cells in which >500 RNA features were expressed and <20% of the genes expressed were mitochondrial. To integrate datasets within stem cell lines, each Seurat object was normalized using the SCTransform pipeline^11^. Seurat objects were then integrated by first finding the integration anchors using the oldest control samples as the reference. For WIBR3 *TSC2^c/-;lslTdTom^* hESCs, the day 240 *TSC2^c/-^* cells were used as a reference. For WIBR3 *TSC2^c/+;LSL-TdTom^* hESCs, the day 120 *TSC2^c/+^* cells were used as a reference, and for BJ *TSC2^c/-;LSL-TdTom^* hiPSCs, the day 140 *TSC2^c/-^* cells were used as a reference. Other parameters were left at their defaults and then the anchor set was integrated. Principal Component Analysis (PCA) was then run on the post-integration cells, followed by UMAP dimensionality reduction using the first 20 PCs. Shared nearest neighbors for each cell and cluster were identified.

The organoid datasets generated in this paper were projected onto a primary fetal tissue dataset^12^ by creating Seurat objects from the publicly available gene-expression matrices, finding the integration anchors between the organoids and the primary tissue data, and then integrating the two datasets into a single Seurat object. Downstream analysis of the integrated dataset was then performed as above. The genes used to calculate module scores are listed in Supplemental Table 2.

Differential expression was performed using MAST, with batch included as a latent variable^13^. Cell type proportions were plotted using the DittoSeq R package^14^, and volcano plots were plotted using the EnhancedVolcano R package^15^. Expression distance analysis was performed using the Cacoa R package^16^. Pathway changes were analyzed using the Single Cell Pathway Analysis (SCPA) R package^17^. Spatial similarity maps were generated using the VoxHunt R package^18^.

### Data availability

The code used to perform all analyses is available as a Github repository [URL Pending]. Raw data is available through Dryad: [DOI Pending].

### Immunostaining of organoid sections

Organoids were fixed with 4% PFA (Electron Microscopy Sciences, 15710) for 2 hours at 4°C. After fixation, organoids were transferred to a 30% sucrose solution and allowed to settle at 4°C overnight or at room temperature for 1 hour. For cryosectioning, organoids were embedded in Tissue-Tek OCT compound (Fisher Healthcare, 4585), frozen in an ethanol and dry ice bath, and sectioned on a cryostat (Leica, CM3050S) into 18 μm sections. Sections were washed once with 1x PBS and blocked in Block-Aid (Thermo Fisher, B10710) with 0.3% Triton X-100 (Sigma, X100) for 1 hour at room temperature. Sections were incubated overnight at 4°C in primary antibodies in Block-Aid. The following day, sections were washed three times with PBS, incubated in secondary antibody (1:500 in Block-Aid) and Hoescht stain (1:1000, Thermo Fisher, H1399) for 1 hour at room temperature and washed again three times with 1x PBS. Slides were coverslipped with ProLong Glass Antifade mountant (Thermo Fisher, P36980) and allowed to cure before imaging. Antibody vendors, catalog numbers, and dilutions are listed in Supplemental Table 13.

2D Immunopanned cultures were fixed with 4% PFA for 15 minutes at room temperature, and immunostained according to the protocol described above, omitting the coverslipping step.

### Confocal imaging and image analysis

All images were acquired using an Olympus Fluoview FV3000 confocal microscope. For organoid sections and 2D cultures, tile scans were collected using a 10X or 20X objective, and stitched using ImageJ’s ‘Grid/Collection Stitching’ plugin using a PyImageJ wrapper. Individual cells were segmented using Stardist^19^ on the nuclei channel, and debris was excluded using an area filter.

For intensity-based measurements, the regionprops_table() function was used with the Stardist labels and the corresponding intensity image. For certain protein targets that were expressed in the cell body but did not include the nucleus, the labels were expanded by 5 pixels to include the soma.

To determine whether a cell was positive or negative for a particular marker, a binarization approach was used (Fig. 3h). The intensity image was post-processed to remove noise or excessive background, and thresholded according to a mean filter. Stardist labels were then overlaid on the resulting binary image, and labels that contained more than 95% positive pixels were considered positive. Each analysis was validated by visualizing positive and negative cells using Napari^20^ to confirm that cells were correctly classified.

In the whole-organoid p-S6 analysis (Fig. 3c), a hybrid approach was used in which intensities from expanded labels were extracted only from cells that contained more than 50% positive pixels in the binarization approach. This approach restricted the analysis only to cells above a minimum detectable level of p-S6, as cells with no detectable p-S6 were likely dead.

In the analysis of standard immunostaining in human tuber samples, only selected regions of tubers were imaged. Cells with fluorescence intensities greater than one standard deviation above the mean in a given image were considered to be high p-S6 cells, with all other cells considered to be low p-S6 cells. In the quantification of cyclic immunofluorescence, the full tuber section was imaged, and the high and low p-S6 thresholds were chosen manually for each tuber due to the variability across samples.

### Western blotting

Brain organoids were harvested in lysis buffer containing 1% SDS in 1× PBS with Halt phosphatase inhibitor cocktail (ThermoFisher: PI78420) and Complete mini EDTA-free protease inhibitor cocktail (Roche: 4693159001). Immunopanned astrocyte cultures were harvested on Day 7 after panning. Neurobasal media was aspirated from the well and wells were quickly rinsed with ice cold 1x DPBS with Ca2+/Mg2+ and then 75 μl of lysis buffer were added (lysis buffer: 10mM Na-PPi (Sigma, 221368), 10mM Na-Beta-glycerophosphate (Sigma, G5422), 40mM HEPES, 4mM EDTA (Sigma, E5134), 1% Triton X-100 (Sigma, T8787), phosphatase inhibitor and protease inhibitor in 1× PBS adjusting pH to 7.4). Total protein was determined by BCA assay (ThermoFisher, PI23227) and 3 or 4 μg of protein in 1X Laemmli sample buffer (Bio-Rad,161-0747) were loaded onto 4–15% Criterion TGX gels (Bio-Rad, 5671084). Proteins were transferred overnight at low voltage to PVDF membranes (Bio-Rad, 1620177), blocked in 5% milk in 1x TBS-Tween for one hour at RT, and incubated with primary antibodies diluted in 5% milk in 1x TBS-Tween overnight at 4°C. The following day, membranes were washed 3 × 10 min in 1x TBS-Tween and incubated with HRP-conjugated secondary antibodies (1:5000) for one hour at RT, washed 6 × 10 min in 1x TBS-Tween, incubated with chemiluminescence substrate (Revvity Health Sciences, NEL105001EA) and developed on GE Amersham Hyperfilm ECL (VWR, 95017-661). Membranes were stripped by two 6 min incubations in stripping buffer (6 M guanidine hydrochloride (Fisher Scientific, ICN10190505) with 1:150 β-mercaptoethanol) with shaking followed by four 2 min washes in 1x TBS with 0.05% NP-40 to re-blot on subsequent days.

Bands were quantified by densitometry using ImageJ 1.52p software (NIH). For all experiments, phospho-proteins were normalized to their respective total proteins. ꞵ-actin was used as a loading control for every experiment. Antibody vendors, catalog numbers, and dilutions are listed in Supplemental Table 6. All antibodies were used in accordance with manufacturer guidelines and were validated by the manufacturer for use in human samples for the specific assays used in this study.

### Immunopanning

Immunopanning was performed on non-treated 6-well plates (Corning, 3736). The day before the experiment, each well was incubated overnight at 4°C with 1:400 goat anti-mouse IgG+IgM antibody (Jackson ImmunoResearch, 115-005-044) in 50 mM Tris-HCl pH9.5 (Thermo Fisher, J62084.K2). Plates for plating cells were prepared by coating Corning BioCoat Poly-D-Lysine 24-well plates (Corning, 356414) with laminin (Sigma, 11243217001) diluted 1:20 in 1x PBS at 37°C overnight.

On the day of immunopanning, panning plates were rinsed 3 times with 1x PBS, then incubated at room temperature with 1:1000 anti-HepaCAM antibody (R&D systems, MAB4108, resuspended at 500 ug/mL) or 1:1200 anti-Thy1 antibody (BD, 550402) in 1 mL of 1x PBS. Plates were incubated until use (∼1 hour). 6-15 organoids were dissociated according to the organoid dissociation for FACS protocol listed above. Organoids were triturated and resuspended in 1 mL of room temperature 0.2% BSA (Sigma, A9418) in 1x PBS with 10 uM Y-27632. After dissociation, cells were filtered with a 70 µm pore size strainer (Greiner, 542170) to eliminate the clumps. Each panning plate was rinsed 5 times with 1x PBS, then the cell suspension was added and incubated at room temperature. First, the cells were incubated on the anti-Thy1 plate for 15 minutes, then the suspension was transferred to the anti-HepaCAM plate for 20 minutes. Loosely bound cells were washed off with additional BSA/PBS/Y27 solution.

After incubation, attached cells were dislodged with 1 mL accutase. Cells on the anti-Thy1 plate were incubated with accutase at room temperature for 10 minutes and cells on the HepaCAM plate were incubated at 37°C for 7 minutes. The accutase was inactivated with 1 mL of neural induction media (see Organoid differentiation section) supplemented with BDNF and NT3, and cells were dislodged using trituration and counted using a hemocytometer.

The laminin was removed from the plating plates, the cell suspension was added to the plate without rinsing, and cells were returned to the incubator for 1 hour. After 1 hour, a full media change was performed with neural induction media supplemented with BDNF and NT3. During this process, centrifugation was avoided whenever possible, as it resulted in substantial sample loss.

### Bulk RNA sequencing

For bulk RNA sequencing, each organoid was transferred to a 1.5 mL tube, residual media was removed, and samples were snap-frozen in liquid nitrogen. RNA was extracted using the RNeasy Mini kit (Qiagen, 74104) according to manufacturer instructions. Library preparation and sequencing was performed by the QB3-Berkeley Genomics core labs. Total RNA quality as well as poly-dT enriched mRNA quality were assessed on an Agilent 2100 Bioanalyzer. Libraries were prepared using the KAPA mRNA Hyper Prep kit (Roche, KK858). Truncated universal stub adapters were ligated to cDNA fragments, which were then extended via 9 cycles of PCR using unique dual indexing primers into full length Illumina adapters. Library quality was checked on an AATI (now Agilent) Fragment Analyzer. Library molarity was measured via quantitative PCR with the KAPA Library Quantification Kit (Roche, KK4824) on a BioRad CFX Connect thermal cycler. Libraries were then pooled by molarity and sequenced on an Illumina NovaSeq X with the 25B flowcell for 2 x 150 cycles, targeting at least 25M reads per sample. Fastq files were generated and demultiplexed using Illumina BCL Convert v4 and default settings.

### Immunostaining of human cortical tuber sections

Cortical tuber samples were obtained from TSC patients undergoing tuber resection surgery, with informed consent under a protocol approved by the Stanford University Institutional Review Board. After surgical resection, samples were placed into Eppendorf tubes and frozen and stored at -80 °C. Portions of samples were cut and embedded in OCT. OCT sample blocks were cryosectioned (Leica, CM3050S) to create 18 μm sections.

For standard immunohistochemistry, cryosectioned samples were fixed with 4% PFA in 1x PBS for 10 minutes and then washed three times in 1x PBS. Sections were blocked in buffer containing 10% normal donkey serum (NDS, Jackson ImmunoResearch, 017-000-121), and 0.3% Triton X-100 in 1x PBS for 1 hour at room temperature. Sections were then incubated overnight at 4°C with primary antibodies in antibody dilution buffer (10% NDS in 1x PBS). The following day, sections were washed three times with 1x PBS, incubated in secondary antibody (1:500 in antibody dilution buffer) and Hoescht stain (1:1000) for 1 hour at room temperature and washed three times with 1x PBS. Slides were coverslipped with ProLong Glass Anti-fade Mountant and allowed to set for at least 1 day before imaging. Antibody vendors, catalog numbers, and dilutions are listed in Supplemental Table 13.

For cyclic imaging, a well was created by cutting a rectangular shape from a sheet of cured polydimethylsiloxane (Dow, Sylgard 184), and pressed to the glass slide with cryosectioned samples. Antibody staining was performed as described above, except that the sample well was filled with an imaging buffer (freshly prepared 0.7 M N-Acetyl-Cysteine (Sigma, A7250) in 0.2 M phosphate buffer at pH 7.4) instead of mounting media^21,22^, and imaging of the full tuber section was performed through the bottom glass slide. Antibody elution was performed immediately after imaging. Prior to elution, TCEP-HCl (Sigma, C4706) was added to a stock solution (0.5 M L-Glycine (Fisher, BP381-1), 3 M Urea (Fisher, U15), and 3 M Guanidine hydrochloride (MP Biomedicals, 101905), stored at 4°C) to a final concentration of 0.07M (20mg/mL). To elute antibodies, samples were rinsed with 1x PBS, incubated for 5 minutes with elution buffer and rinsed with water. This process was repeated for a total of 3 washes. After elution, samples were washed with 1x PBS, stained with Hoescht, and selected regions were re-imaged to ensure that elution was successful (Extended Data Fig. 7b). The blocking step for the next round of antibody staining was started immediately afterward. Eluted samples could be stored in the dark at 4°C in 1x PBS for at least 48 hours between cycles, and stained samples containing fluorescent antibodies could be stored in the dark at 4°C in imaging buffer for at least 24 hours.

### Cyclic Staining Analysis

Cyclic staining images were acquired as described above, using a 10X objective and 4096×4096 pixel resolution on an Olympus Fluoview FV3000 microscope. Each cycle was stitched and aligned on the Hoescht channel using a developmental branch of the Alignment by Simultaneous Harmonization of Layer/Adjacency Registration (ASHLAR) software package^23^ containing a rotation correction^24^. Standard intensity-based image analysis was performed as described above, using manually chosen p-S6 intensity thresholds.

The image UMAP process is described in Extended Data Fig. 7a. Beginning with the ASHLAR-aligned image channels, Stardist was used to segment nuclei for each cycle individually. Because ASHLAR alignment results in the same coordinate system for all channels, each nucleus label in the first channel was matched to the nearest label in each subsequent channel to identify the same cell across channels. Poor matches were filtered by a defined maximum allowable distance between any pair of label centroids across channels. After identifying confidently aligned cells, intensity data for each cell was extracted. All pixels within a 30px (9.3 μm) radius circle centered on each cycle’s label centroid across all cycles were extracted. This data was then reshaped into a 1-D vector, and data from all cells was stacked to generate a 2-D matrix of [cell index] X [pixel intensity]. The UMAP-learn package was used to perform dimensionality reduction on this matrix, and HDBSCAN was used on the UMAP embeddings to determine clustering. For differential intensity measurements, intensity data, cluster assignments, and UMAP embeddings were used to generate a Seurat object, and differential intensity was calculated using the Wilcox rank-sums test with the Bonferroni correction.

